# Bortezomib abrogates temozolomide-induced autophagic flux through an ATG5 dependent pathway

**DOI:** 10.1101/2020.12.20.423718

**Authors:** Mohummad Aminur Rahman, Agnete Engelsen, Shahin Sarowar, Christian Bindesbøll, Even Birkeland, Maria L. Lotsberg, Stian Knappskog, Anne Simonsen, Martha Chekenya

## Abstract

Glioblastoma (GBM) is invariably resistant to temozolomide (TMZ) chemotherapy. Inhibiting the proteasomal pathway is an emerging strategy to accumulate damaged proteins and inhibit their lysosomal degradation. We hypothesized that bortezomib (BTZ) might sensitize GBM cells to TMZ. We examined change in autophagic flux after drug treatments and in combination with pharmacological inhibitors or CRISPR cas9 knockout of autophagy-related genes -5 and -7 (ATG5 and ATG7, respectively). Autophagic flux was increased in temozolomide resistant GBM cells as indicated by diminished levels of the autophagy markers LC3A/B-II and p62(SQSTM1), increased localisation of LC3A/B-II with STX17, higher long-lived protein degradation and no induction of apoptosis. In contrast, BTZ treatment abrogated autophagic flux by accumulation of LC3A/B-II and p62(SQSTM1) positive autophagosomes that did not fuse with lysosomes and reduced degradation of long-lived proteins. BTZ synergistically enhanced TMZ efficacy by attenuating cell proliferation, increased DNA damage and apoptosis. CRISPR Cas *ATG5* knockout reversed BTZ-induced autophagy blockade and rescued the GBM treated cells from death. We conclude that bortezomib abrogates temozolomide induced autophagy through ATG5 dependent pathway.

## Introduction

Glioblastoma (GBM) is the most frequent, lethal primary brain malignancy in adults and is inherently chemoresistant. Standard therapy includes surgery, 75 mg/m^2^ temozolomide (TMZ) alkylating chemotherapy administered concomitantly with 2-Gy fractionated ionizing radiotherapy daily for 5 days in six consecutive cycles (chemoradiation with 60-Gy total radiation dose). Thereafter, patients continue with a 5-day treatment cycle of 150 mg/m^2^ TMZ for the first month, followed by 5 cycles of 200 mg/m^2^ for 5-days every month if well tolerated. Despite this aggressive schedule, median survival is only 14.6 months^1^. Thus, there is an urgent, unmet need for strategies that can overcome chemoresistance as society ages and GBM incidence might be expected to increase.

The cytotoxic effect of TMZ is mostly driven by the O^6^-methylguanine adducts that cause a preferential mismatch pairing of guanine with thymine rather than cytosine leading to genomic instability. The generated single- and double-strand DNA breaks ultimately trigger tumor cell death by apoptosis^2^. However, a major mechanism that counteracts this cytotoxicity, is the DNA repair enzyme, O^6^-methylguanine DNA methyltransferase (MGMT) which sequesters the toxic methyl-adducts from O^6^ guanine prior to DNA replication^3,4^, rendering the tumor cells resistant to TMZ cytotoxicity. Consequently, gene silencing by promoter methylation of the *MGMT* gene is a strong prognostic and predictive factor for response to TMZ^5^. However, *MGMT* promoter hypomethylation is not the only mechanism for chemoresistance, as GBMs also harbor mutations in base excision repair genes^6,7^ that may underlie the heterogeneity in patients’ responses to TMZ. This may explain why some tumors are resistant to TMZ despite methylated *MGMT* promoter. Moreover, recurrent tumors may acquire a chemo-resistant phenotype denoted by whole genome enrichment of the C:G>T mutational signatures. Further gain of inactivating mutations in components of the mismatch repair pathway^8^ is also indicative of TMZ-induced mutagenesis.

Notwithstanding genetic alterations that promote drug resistance, a common survival strategy for many cancers is metabolic reprogramming characterized by increased macroautophagy^9,10^ (hitherto referred to as autophagy), a well conserved catabolic process that allows sequestration of long-lived cellular proteins, toxic protein aggregates and damaged organelles into double-membrane autophagosomes. The cargo is subsequently degraded after fusion of the autophagosomes with lysosomes^11,12^, to form autolysosomes. This process will henceforth be referred to as autophagic flux. The formation of autophagosomes is a multistep process involving several proteins, including the serine/threonine Unc-51-like Autophagy-Activating-Kinases 1 and 2 (ULK1 and ULK2), forming a multi-subunit ULK-complex, which initiates the formation of the phagophore membrane^13,14,15^ by recruiting the Vacuolar Protein Sorting-associated protein (VPS34, a class III phosphatidyl ionositol-3 kinase) and downstream proteins that further promote elongation and closure of the phagophore to form an autophagosome^16^. Protein products of several autophagy-related genes (ATG), including ATG5 and ATG7, mediate conjugation of the ubiquitin-like microtubule-associated protein 1A/1B-light chain 3 (MAP1LC3A/B, hereafter referred to as LC3), to phosphatidylethanolamine in autophagic membranes (referred to as LC3A/B-II). LC3A/B-II can interact with autophagy receptors e.g p62/Sequestosome-1 (p62/SQSTM1), which further recruit ubiquitin-tagged cargo into autophagosomes. Fusion of the latter with acid hydrolase containing lysosomes is controlled by SNARE proteins, including syntaxin 17 (STX17), resulting in formation of autolysosomes where cargo becomes degraded into components that can be reused by the cell^14^.

Autophagic flux is thus postulated to function as an adaptive survival mechanism that sustains cancer cells during conditions of stress^17–20^. Transcriptional regulation by members of the forkhead homeobox-type O (FOXO), as well as deregulated signaling pathways involving the MEK/ERK and PI3K are known to affect basal rates and dependence on autophagic flux in different conditions and cancer types^10,15,21,22^. Indeed, increased autophagic flux after TMZ treatment has been proposed to represent an additional mechanism for TMZ resistance in GBM cells^23,24^. As a result, combination of anti-cancer treatments with FDA approved inhibitors of autophagy flux, such as chloroquine or hydroxychloroquine have been tested in clinical trials^25^ to potentiate therapeutic efficacy of TMZ^23,26,22^.

Cancer cells may nevertheless acquire resistance to death by abrogated autophagy, serendipitously rendering them more susceptible to proteasome inhibition^27^. Inhibiting the 26S proteasome has emerged as an attractive strategy to accumulate damaged or misfolded proteins, reactive oxygen species and endoplasmic reticulum stress, ultimately lowering the apoptosis threshold of tumor cells^28,29^. New generation reversible and non-reversible proteasome inhibitors, such as Ixazomib and Marizomib (respectively) are currently under clinical investigation in combination with anti-neoplastic treatments in cancers, including GBM. Like Ixazomib, Bortezomib (BTZ, Velcade) reversibly blocks the chymotryptic-like activity of the β1 and β5 subunits of the 26S proteasome. BTZ is approved for treatment of multiple myeloma and mantle cell lymphoma and has been trialed in early phase clinical studies for GBM^31–34^. *In vitro* studies with BTZ alone or in combination with other drugs showed potent anti-cancer activity against various tumors through multiple mechanisms^35,36^. We demonstrate here that pre-treatment with BTZ prior to TMZ potently sensitized GBM cells to TMZ, as indicated by reduced IC_50_ doses, rapidly abrogated autophagic flux and strongly induced DNA damage response and apoptosis in an ATG5 dependent manner.

## Results

### Bortezomib synergistically enhances sensitivity of GBM cells to temozolomide chemotherapy

We first investigated the clonogenic survival of a panel of patient-derived GBM cells and cell lines in response to various doses of TMZ and BTZ alone or in combination. P3, T98G and HF66 tumor cells were most resistant to TMZ, requiring higher doses (>50μM) to kill 50% of the cells, compared to U87 and A172 tumor cells whose IC_50_ doses were 12 and 25μM, respectively (*P*=0.0001, Fig. 1A). In contrast, all tumor cell lines, as well as control cells (immortalized normal human astrocytes (NHA)) were sensitive to low doses of BTZ (IC_50_ range 6-12nM, Fig. 1B). Pretreatment with 5nM, 10nM and 15nM BTZ prior to TMZ effectively reduced the clonogenic survival of P3 and T98G cells compared to TMZ treatment alone. BTZ pretreatment potentiated the effect of all TMZ doses for both P3 and T98G cells (*P*<0.0001, Fig. 1C and 1D, respectively), where 15nM BTZ was synergistic with all TMZ doses (Fig. 1E). Taken together, pretreatment with BTZ rescinded GBM cells’ resistance to TMZ.

**Figure 1.**
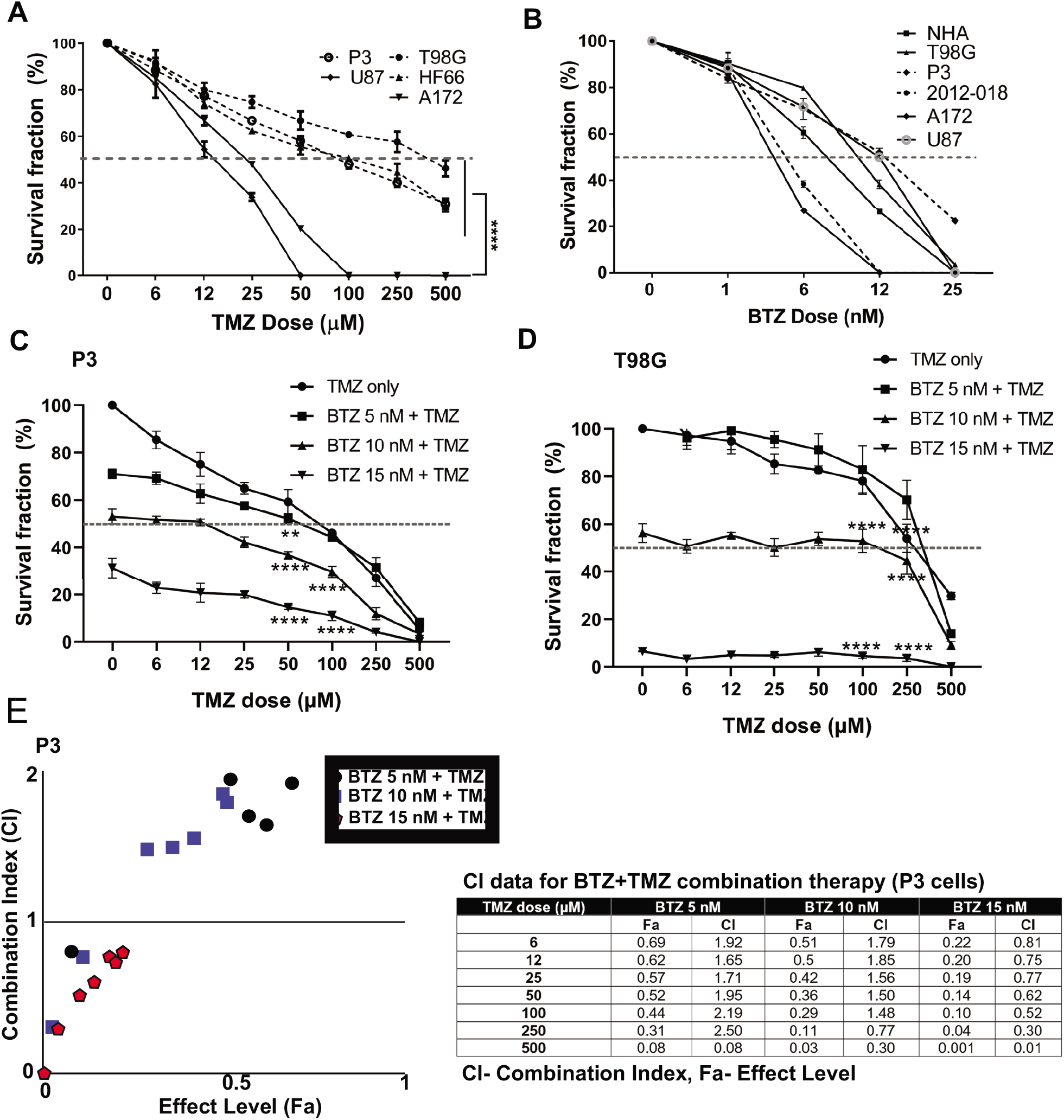
BTZ pretreatment synergizes with TMZ treatment and chemosensitizes GBM cells. (A) Mean % ± S.E.M clonogenic survival of tumor cells treated with TMZ for 72 h (B) Mean % ± S.E.M clonogenic survival of tumor cells and normal human astrocytes (NHA) treated with BTZ for 48h. Clonogenic surviving fractions of (C) P3 and (D) T98G tumor cells after specified treatments, BTZ concentration was 0, 5, 10 and 15nM, respectively, and TMZ concentration in 6-500 μM range. (E) Combination index plot (left) and data table (right) from report generated by Compusyn software for drug combination synergy calculation following the Chou-Talalay method of P3 cells treated with BTZ 5, 10 and 15nM and TMZ concentration in 6-500μM range. Each experiment was performed in triplicate, and data represents the mean ± S.E.M of at least 3 independent experiments, * P < 0.05, ** P < 0.01, *** P < 0.001 and **** P < 0.0001.

### Genetic background of GBM cells

To rule out that differences in genetic mutations accounted solely for the differential responses to BTZ and TMZ treatment, we characterized the genetic background of P3 and T98G cells, Table 1. Both cell types harbored wild-type (Wt) isocitrate dehydrogenase 1 gene (*IDH1*) that denoted them as *de novo* GBM. P3 cells harbored a homozygous *Cys176Tyr* nonsynonymous single nucleotide variant (SNV) in the tumor protein 53 (*TP53*) gene, whilst T98G cells possessed a *Met237Ile* SNV mutation in *TP53*. Both P3 and T98G cells had mutations in *MAP3K1* and in the phosphatase tensin homolog (*PTEN*) gene, while additionally, P3 cells harbored a heterozygous *PIK3R2* mutation in exon 11 and in *STAT3*, potentially inducing aberrant PI3K signaling (Table 1 and supplementary Table 2). Particularly in P3 cells, these aberrations are implicated in development of autophagy addiction^37^.

**Table 1:**
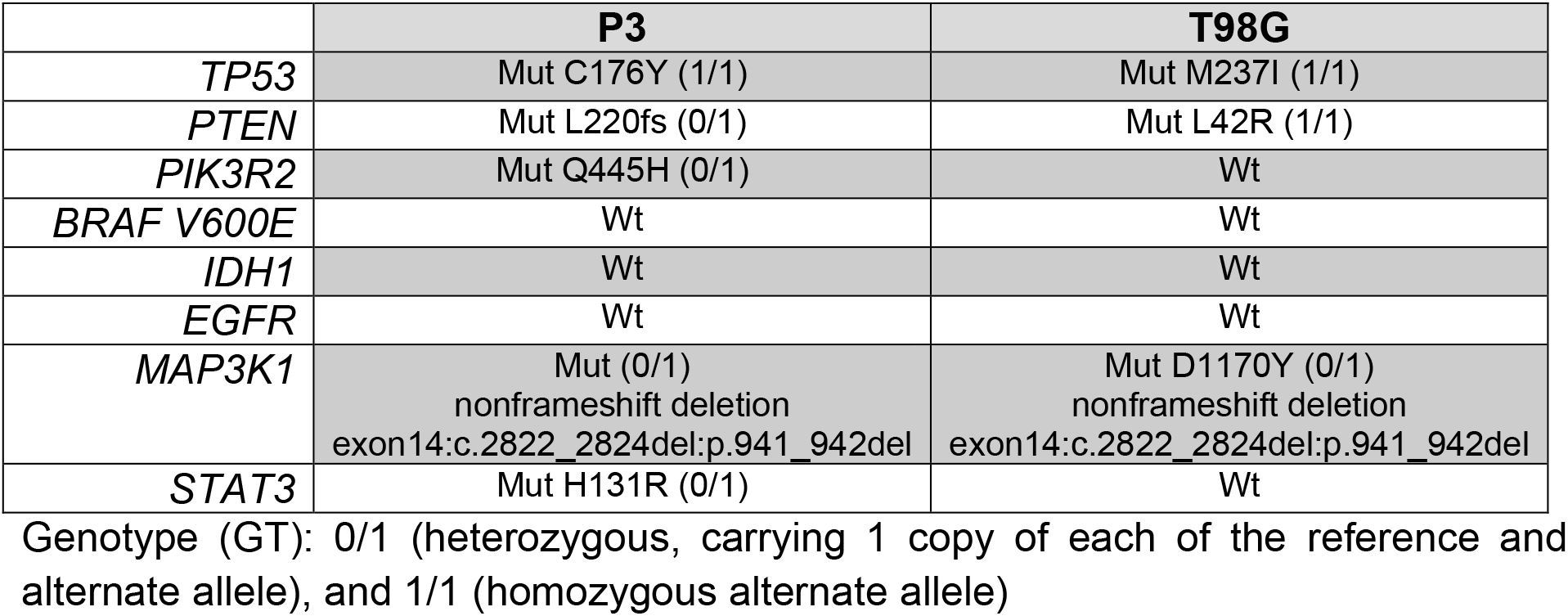
Genetic background of GBM cell lines

|  | **P3** | **T98G** |
| --- | --- | --- |
| *TP53* | Mut C176Y (1/1) | Mut M237I (1/1) |
| *PTEN* | Mut L220fs (0/1) | Mut L42R (1/1) |
| *PIK3R2* | Mut Q445H (0/1) | Wt |
| *BRAF V600E* | Wt | Wt |
| *IDH1* | Wt | Wt |
| *EGFR* | Wt | Wt |
| *MAP3K1* | Mut (0/1)<br>nonframeshift deletion<br>exon14:c.2822_2824del:p.941_942del | Mut D1170Y (0/1)<br>nonframeshift deletion<br>exon14:c.2822_2824del:p.941_942del |
| *STAT3* | Mut H131R (0/1) | Wt |

Genotype (GT): 0/1 (heterozygous, carrying 1 copy of each of the reference and alternate allele), and 1/1 (homozygous alternate allele)

### BTZ and BTZ+TMZ combination treatment induce an accumulation of lipidated LC3A/B (LC3A/B-II) and p62(SQSTM1) in patient derived GBM cells

Autophagy is frequently deregulated in cancer and is considered a maladaptive process, promoting both tumor cell survival or death at different stages of disease progression or in response to treatment^23,38^, in a cell specific manner and in response to proteasome inhibitors. Since TMZ monotherapy was unable to efficiently kill GBM cells, we hypothesized that increased autophagic flux might have sustained survival of GBM cells during TMZ treatment. The levels of the autophagy marker LC3A/B-II in protein lysates from P3 cells after TMZ treatment were minimally altered relative to untreated controls, whereas the level of the autophagic cargo receptor p62(SQSTM1) was diminished by approximately 40% (Fig. 2A), potentially indicating sustained autophagy. In contrast, treatment with 10nM BTZ induced accumulation of both LC3A/B-II and p62(SQSTM1) protein levels by approximately 2-fold after 48h (Fig. 2A). However, 15nM BTZ increased LC3A/B-II and p62(SQSTM1) protein levels by 14 and 8-fold, respectively, relative to untreated controls (Supplementary Fig. 1). Combining 10nM BTZ with different TMZ doses and time points resulted in increased levels of LC3A/B-II (approximately 4-fold) and p62(SQSTM1) levels approximately 2-fold (Supplementary Fig. 1), potentially indicating accumulated autophagosomes. In T98G cells, LC3A/B-II levels were only induced by combination 10nM BTZ + 250μM TMZ as well as chloroquine (CQ) treatment with modest changes in p62(SQSTM1) (Supplementary Fig. 2A) potentially indicating reduced susceptibility in autophagosome accumulation in these cells.

**Figure 2.**
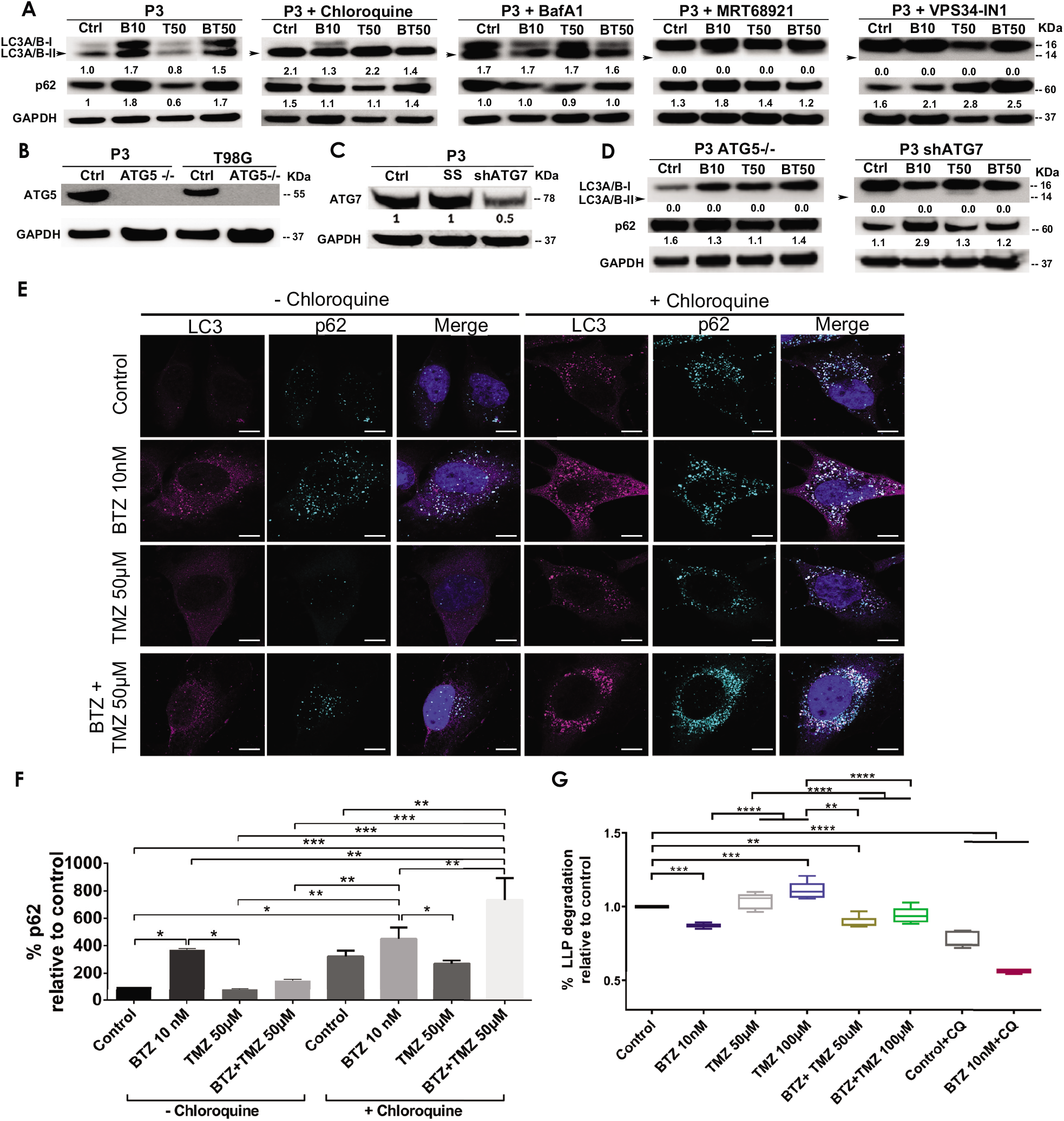
BTZ and combination BTZ+TMZ treatment accumulates LC3A/B-II and p62(SQSTM1) in patient derived GBM cells. (A) Western blot of the indicated proteins in lysates of wild type P3 cells and P3 cells treated with 10nM BTZ (B10), 50μM TMZ (T50) or both (BT50) or in a combination with autophagy inhibitors (chloroquine, Bafilomycin A1 (BafA1), MRT68921 or VPS34-IN1). (B-C) Western blot of the indicated proteins in lysates of wild type P3, T98G cells and cells lacking *ATG5* (*ATG5^-/-^*), scrambled shRNA (SS) and ATG7 (shATG7). (D) Western blot of the indicated proteins in lysates of wild type P3 cells lacking *ATG5* (*ATG5^-/-^*) or ATG7 (shATG7), treated with 10nM BTZ (B10), 50μM TMZ (T50) or both (BT50). GAPDH was used as a loading control and for normalizing densitometry measurements of the blots. (E) Confocal microscopy of P3 cells stained with antibodies against LC3A/B and p62(SQSTM1) after the indicated treatments in the absence (left panel) or presence (right panel) of chloroquine (CQ), scale bar 10μm. (F) Quantification of p62(SQSTM1) puncta after the indicated treatments in the absence or presence of CQ autophagy inhibition. Data represents % mean ± S.E.M from at least 10 cells from 2-3 independent experiments. (G) Percent degradation of long-lived proteins (LLPD) relative to untreated control, quantified as release of ^14^C-valine after indicated treatments in the presence or absence of CQ. * P < 0.05, **P < 0.01, *** P < 0.001 and **** P < 0.0001.

### BTZ and BTZ+TMZ mediated accumulation of LC3A/B-II and p62(SQSTM1) autophagosomes in GBM cells is dependent on core autophagy machinery

To investigate whether the increase in LC3A/B-II and p62(SQSTM1) protein levels in BTZ as well as BTZ+TMZ treated cells was autophagy-specific and identify the particular stage affected in the multistep process, we targeted ULK1/2 and PIK3C3/VPS34 proteins that are both involved in early phagophore initiation. Both MRT68921 and VPS34-IN1, inhibitors of ULK1/2 and VPS34 respectively, prevented LC3A/B lipidation (Fig. 2A). In contrast, 2-3 fold elevated p62(SQSTM1) levels were detected in cells co-treated with BTZ and these inhibitors. The late stage autophagy inhibitors Bafilomycin A1 (BafA1) and CQ both inhibit autophagosome-lysosome fusion by affecting acidification^39^, and like BTZ, they also induced approximately 2-fold elevation of LC3A/B-II levels in the absence, but *not* presence of BTZ (Fig. 2A). Elongation and expansion of the phagophore is enabled by conjugation of LC3A/B-II to phosphatidylethanolamine (PE), facilitated by the E1 ligase ATG7 and the E3-like complex ATG12-ATG5-ATG16L1. Targeting *ATG5* in this conjugation complex by CRISPR-Cas9 completely abolished ATG5 protein (Fig. 2B) whereas small hairpin RNA mediated knockdown of ATG7 mRNA (ATG7 shRNA) depleted 50% of the protein (Fig. 2C). Both *ATG5^-/-^* and ATG7 shRNA cells completely lacked lipidated LC3A/B and the p62(SQSTM1) level was upregulated by approximately 2-fold in *ATG5^-/-^* cells, but not after BTZ treatment (Fig. 2D). In contrast, BTZ treated ATG7 shRNA cells were still able to upregulate p62(SQSTM1) by approximately 3-fold (Fig. 2D).

In line with these western blot data, LC3A/B and p62(SQSTM1) puncta increased after treating P3 cells with BTZ alone or in combination with TMZ, but not under steady state or when treated with TMZ alone (*P* < 0.0001, *P* < 0.05 and *P* > 0.05, respectively, Fig. 2E and 2F). p62(SQSTM1) positive puncta were also prominently observed in the presence of the lysosomal inhibitor CQ in P3 cells (Fig. 2E and 2F) suggesting that BTZ accumulates autophagosomes which may not degrade, as does the late-stage inhibitors, BafA1 and CQ.

### Increased autophagosome accumulation after BTZ and BTZ+TMZ treatment is due to block in autophagic flux as a result of diminished autophagosome-lysosome fusion

To confirm that increased autophagosome formation after BTZ treatment was due to abrogated cargo degradation and not increased autophagic flux, we assessed the effect of the treatment on long-lived protein degradation (LLPD) of the treated cells. Cells were pulsed with radiolabeled valine and following a chase to deplete short lived proteins and induction of autophagy by starvation, the degradation of long-lived proteins, which is mainly mediated by autophagy, was quantified (Fig. 2G). Indeed, the level of recycled amino acids was higher after treatment with 100μM TMZ in P3 cells compared to control (*P* < 0.001) and in TMZ 100μM *vs*. BTZ+TMZ 100μM (*P* <0.0001, Fig. 2G). In contrast, BTZ monotherapy significantly reduced the degradation of long-lived proteins compared to untreated control (*P* < 0.0001) and both TMZ 50μM and 100μM (*P* <0.0001, Fig. 2G), consistent with an abrogated cargo degradation, *ergo* abrogated autophagic flux. BTZ+TMZ 50μM also attenuated the degradation of long-lived proteins compared to controls and TMZ 50μM (*P* < 0.01 respectively, Fig. 2G) and confirmed the inhibition of autophagic flux by blocking degradation of cargo. As expected, the autophagy inhibitor CQ inhibited autophagic flux in all treatment conditions (*P* < 0.0001, Fig. 2G), similarly to BTZ treatments. These findings were corroborated in T98G cells, where BTZ containing treatment regimen also attenuated degradation of long-lived proteins compared to untreated controls or TMZ chemotherapy (*P* < 0.0001, Supplementary Fig. 2B).

As a further confirmation that BTZ alone and in BTZ+TMZ combination treatment abrogated autophagic flux, we used the Autophagy Tandem Sensor RFP-GFP-LC3B probe to dynamically monitor the formation of autophagosomes *vs* autolysosomes over time in the presence or absence of CQ. Both untreated controls and TMZ monotherapy treated cells predominantly displayed red-only puncta, representing degradative autolysosomes, indicating that autophagic flux was retained during TMZ treatment (Fig. 3A and 3B). In contrast, BTZ treatment alone or BTZ+TMZ combination treatment blocked autophagic flux, as indicated by a 60-70% increase of yellow puncta (ratio of green *vs* red puncta) autophagosomes relative to untreated control cells (*P*<0.05) as well as BTZ alone compared to TMZ 50μM (*P* < 0.01); TMZ 100μM (P<0.001) or BTZ+TMZ 50μM and BTZ+TMZ 100μM, *P*<0.001 for both, Fig. 3A and 3B). As expected, addition of CQ increased the frequency of yellow puncta, representing autophagosomes, under all treatment conditions (Fig. 3A and 3B).

**Figure 3.**
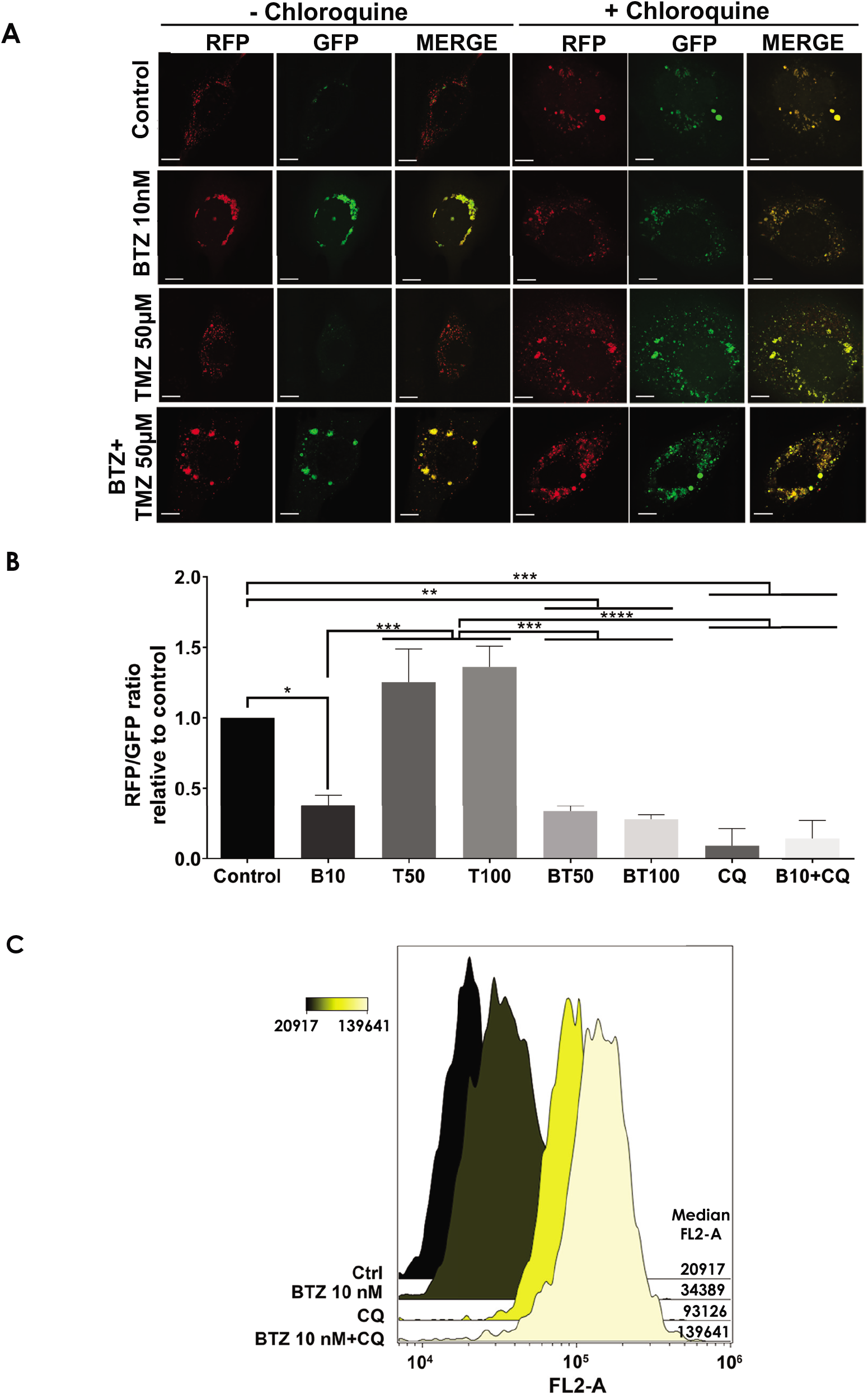
LC3A/B+ autophagosomes accumulate in response to BTZ and combination BTZ+TMZ treatment of GBM cells. (A) Confocal microscopy of P3 cells transfected with RFP-GFP-LC3B tandem probe and subsequently treated as indicated, red (RFP) fluorescence marks autolysosomes and yellow (merge of RFP and GFP) autophagosome puncta formation in the presence or absence of CQ, scale bar 10μm. (B) Quantification of RFP puncta relative to GFP puncta in cells treated with 10nM BTZ (B10), 50μM TMZ (T50), 100μM TMZ (T100) or both (BT50 and BT100) in the absence or presence of CQ, normalized to untreated control cells. * P < 0.05, **P < 0.01, *** P < 0.001 and **** P < 0.0001. (C) Autophagic flux under 3h starvation in EBSS medium of P3 cells was assessed by CytoID probe following pre-treatment as indicated (BTZ 10nM 48h, Chloroquine (CQ) 50μM 24h). Histograms are representative of three independent experiments. Color scale represents median fluorescence intensity (MFI) and values for each treatment are shown on the right side of the histograms.

Autophagic flux at the single-cell level was quantified by flow cytometry using the Cyto-ID probe, which stains autophagic vesicles. When autophagic flux was abrogated by BTZ treatment in nutrient-deprived cells, relative median fluorescence intensity was increased by approximately 2-fold relative to untreated controls (Fig. 3C). Likewise, treatment with CQ strongly induced the accumulation of autophagic vesicles, as indicated by 4-fold increase in median fluorescence intensity relative to control cells. Addition of CQ to the BTZ pre-treated cells had an additive effect and approximately 6-fold increase in median fluorescence intensity (Fig. 3C). Taken together, these data indicate that BTZ treatment abrogates autophagic flux by blocking fusion of autophagosomes with lysosomes under both nutrient-rich and nutrient-deprived conditions.

### BTZ abrogates autophagic flux by blocking autophagosome-lysosome fusion in an ATG5 dependent manner

Confusion exists regarding whether proteasome inhibitors augment^40,41^ or prohibit autophagic flux^42,43^, largely due to lack of use of proper flux assays and misinterpretations of LC3A/B immunoblotting as a surrogate marker for flux^44^. As proof of concept, we targeted the key autophagy regulator proteins involved in the five stages of autophagy: phagophore initiation/nucleation (UKL1/2 and VPS34), expansion and elongation (ATG5 and ATG7), autophagosome closure and lysosomal fusion (bafilomycin A1 and chloroquine) and examined changes in expression of ULK1/2, STX17, LC3A/B-II, and p62(SQSTM1) before and after 48h treatment with BTZ by immunofluorescence, subcellular morphology, western blot of cell lysates, as well as cell clonogenic survival (Fig. 4 and 5).

**Figure 4.**
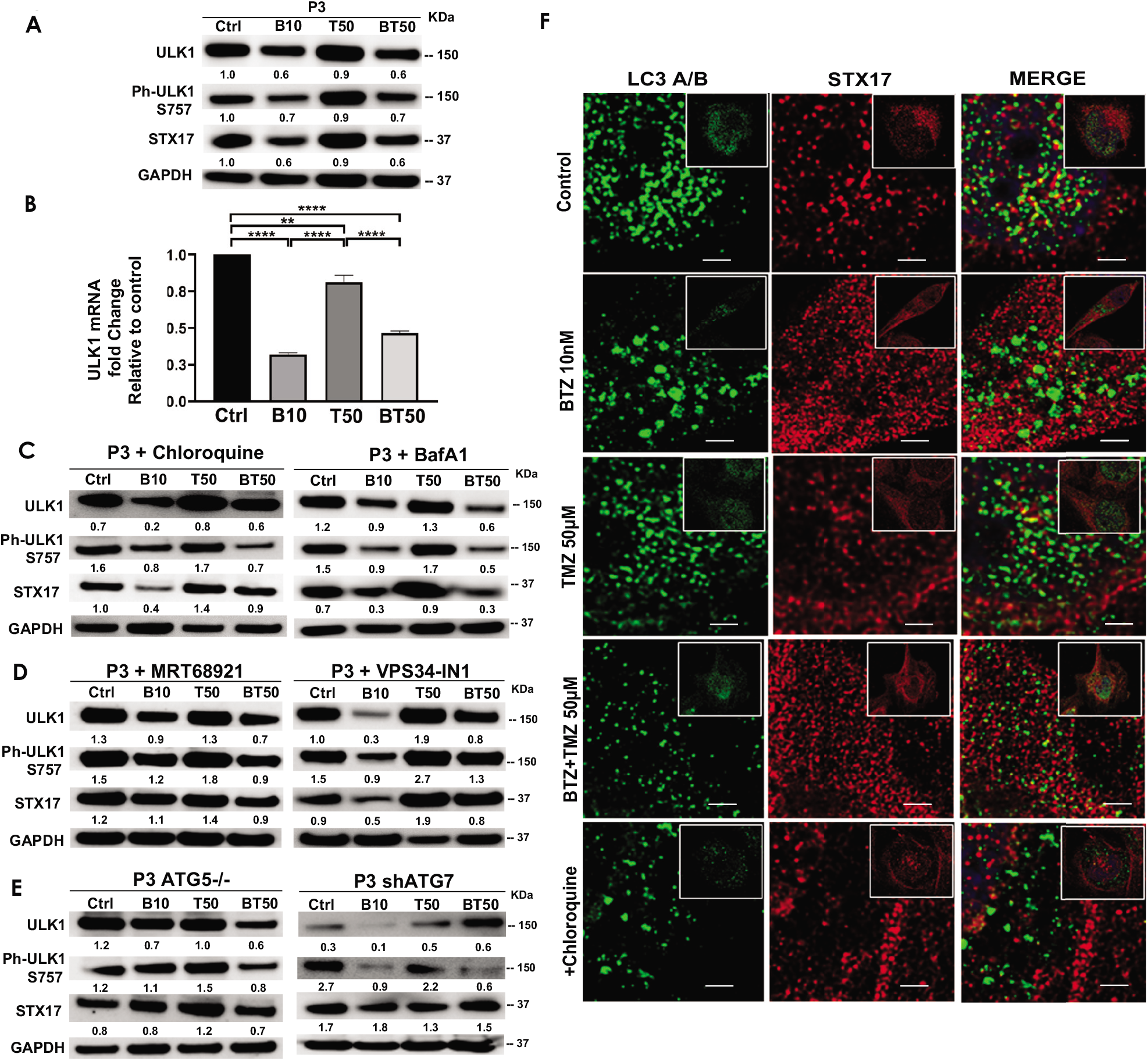
BTZ abrogates autophagic flux by blocking autophagosome-lysosome fusion. (A) Western blot of the indicated proteins in lysates of wild type P3 cells treated with 10nM BTZ (B10), 50μM TMZ (T50) or both (BT50) (B) ULK1 mRNA fold change after indicated treatments in P3 cells. (C-D) Western blot of the indicated proteins in lysates of P3 cells treated with 10nM BTZ (B10), 50μM TMZ (T50) or both (BT50) or in a combination with autophagy inhibitors (chloroquine, BafA1, MRT68921 or VPS34-IN1) (E) Western blot of the indicated proteins in lysates of P3 cells lacking *ATG5* (*ATG5^-/-^*) and ATG7 (shATG7). GAPDH was used as a loading control and for normalizing densitometry measurements of the blots. (D) Confocal microscopy of P3 cells co-stained for LC3A/B and STX17, subsequently treated as indicated, in the presence or absence of CQ, scale bar 3μm; for the zoomed part of the image from the image shown in the white square.

**Figure 5.**
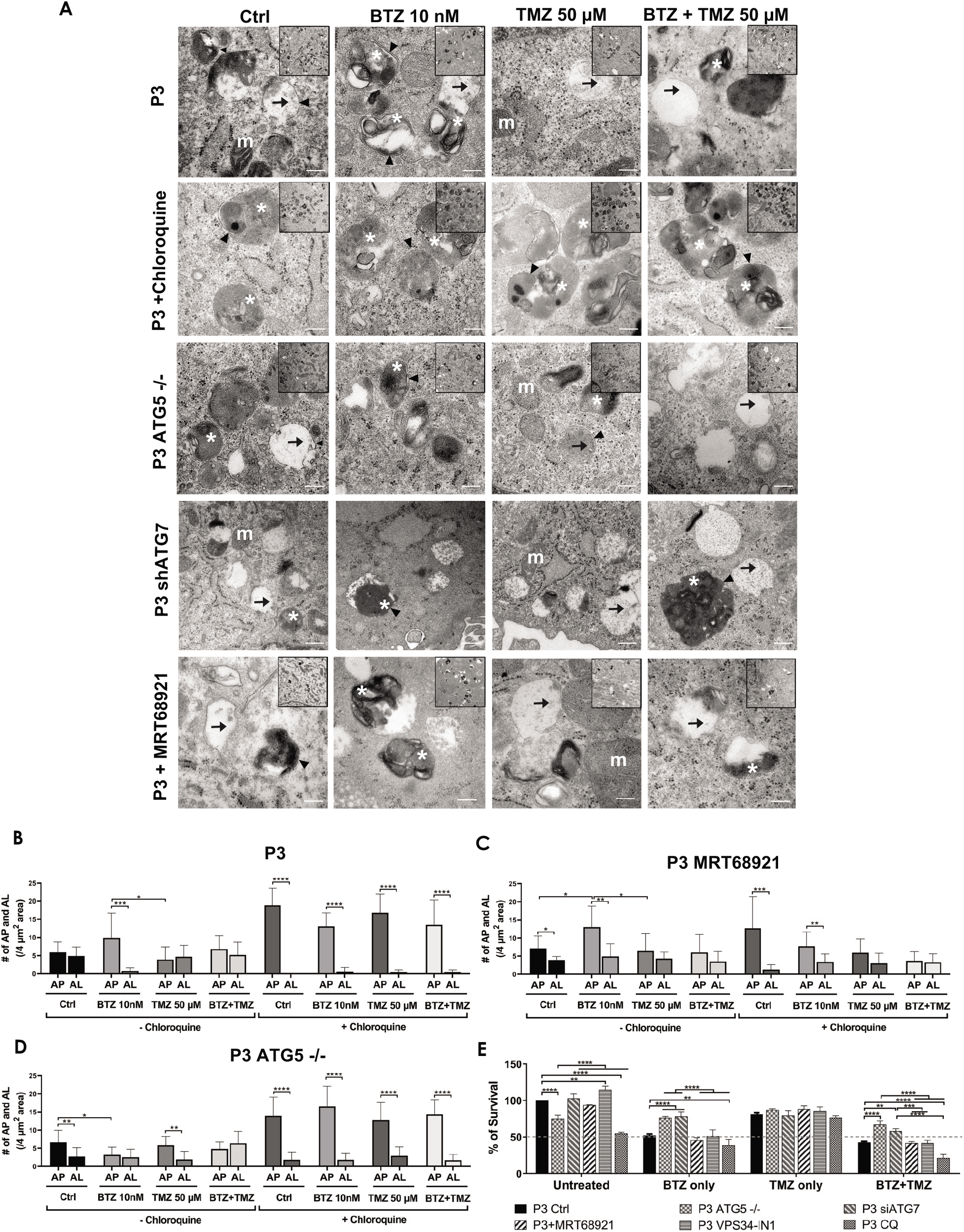
Targeting autophagy diminishes tumor cell survival. (A) Transmission electron microscopy of untreated, BTZ or TMZ monotherapy and BTZ+TMZ combination treated P3 ctrl cells, *ATG5^-/-^* P3 cells and ATG7 knockdown P3 cells as well as P3 cells treated with chloroquine and MRT68921, scale bar 200 nm, magnification 50K. Inserts 20K magnification. Arrowhead=double membrane, white asterix= autophagosomes, arrow=autolysosomes and m (white)=mitochondria. (C-D) Number of autophagosomes and autolysosomes were counted from at least 10 cells from 2-3 independent experiments by two independent investigators in P3 control cells, P3 cells lacking *ATG5* (*ATG5^-/-^*) and P3 cells treated with MRT68921 with or without CQ. (E) Clonogenic survival of untreated or treated P3 control, *ATG5^-/-^*, shATG7, MRT68921, VPS34-IN1and CQ treated cells after experimental treatments. \**P* < 0.05, ** *P* < 0.01, *** *P* < 0.001 and **** *P* < 0.0001.

P3 GBM cells treated with BTZ or BTZ+TMZ had 30-40% attenuated total and phosphorylated ULK1 expression (Fig. 4A), which was *not* restored in the presence of CQ, and as reflected in 60-70% reduction in mRNA levels compared to control and TMZ treated cells (*P*<0.0001, respectively, Fig. 4B). In contrast, ULK1 levels in TMZ treated cells were more similar to untreated controls (*P*<0.01). BTZ and BTZ+TMZ treated P3 cells diminished expression of STX17 marker for lysosomes by 40-60% in presence or not of CQ (Fig. 4C). Targeting ULK1 and VPS34 protein complexes involved in phagophore nucleation with MRT68921 (UKL1/2 dual inhibitor) and VPS34-IN1 depleted lipidated LC3A/B regardless of treatment (Fig. 2A), although BTZ 10nM in presence of VPS34-IN1, STX17 expression was further reduced by 50% (Fig. 4D). Targeting *ATG5* during BTZ treatment had no major impact on phospho-ULK1 or STX17 expression, but ATG7 shRNA cells treated with TMZ increased phospho-ULK1 and STX17 by 2fold (Fig. 4E). Immunofluorescence staining for STX17 and LC3A/B on treated cells revealed absence of STX17/LC3A/B colocalization in BTZ or CQ treated cells but colocalization was evident in untreated controls and TMZ treated cells, (Fig. 4F). Taken together, the data indicate that BTZ depletes ULK1 phosphorylation and accumulates LC3A/B-II in an *ATG5* dependent manner in the early stages, subsequently abrogating flux by blocking autophagosome-lysosome fusion, indicated by low levels of STX17 colocalization with LC3A/B.

Transmission electron microscopy confirmed the equal presence of autophagosomes as well as fused degradative autolysosomes in control and TMZ 50μM treated cells (Fig. 5A and 5B), but 5-fold increase in double-membrane limited autophagosomes containing preserved organelles in BTZ treated cells compared to autolysosomes (*P* <0.001, Fig. 5B) and compared to TMZ 50μM treated cells (*P* <0.05). As expected, autophagosomes were significantly increased compared to autolysosomes regardless of treatment (*P* <0.0001) in the presence of CQ. Corroborating the observed increases in lipidated LC3A/B, inhibition of ULK1/2 with MTR68921 in control and BTZ treated cells significantly increased the frequency of autophagosomes relative to autolysosomes (*P* <0.05 and *P* <0.01, respectively), that were also *not* further augmented in the presence of CQ (*P* <0.001 and *P* <0.01, respectively), and compared to control and TMZ 50μM treated cells, (*P* <0.05 respectively, Fig. 5A and 5C). In the *ATG5* knockout cells, only control and TMZ treated cells displayed an increased ratio of autophagosomes relative to autolysosomes, (*P* <0.01), wherein these *ATG5* cells, BTZ treatment failed to induce autophagosomes (*P* <0.05, Fig. 5A and 5D). As expected in the presence of CQ autophagosomes were increased in all treatment conditions (*P* <0.0001), and when effect of BTZ was abrogated by CRISPR-Cas9 *ATG5^-/-^* (Fig. 5A and 5D).

### Diminished tumor cell survival due to BTZ abrogated autophagic flux requires a functional autophagy machinery

To be certain that BTZ mediated abrogation of autophagic flux was responsible for the biological effects on GBM cell death during combination treatment with TMZ, colony formation as an indication of clonogenic survival was monitored in P3 and T98G cells depleted of *ATG5* or ATG7, or treated with MRT68921 (ULK1/2 inhibitor) or VPS34-IN1 (VPS34 inhibitor) under steady state and during BTZ and TMZ treatment. CRISPR-Cas9 ablation of *ATG5* in both P3 and T98G cells reduced clonogenic survival compared to wild type cells (*P* < 0.0001 and *P* < 0.01, respectively) and compared to wild type cells treated with MRT68921, VPS34-IN1 to inhibit phagophore initiation (early step), or chloroquine to inhibit lysosomal acidification (late step) (*P* < 0.0001, Fig. 5E and Supplementary Fig. 2C). ATG7 shRNA did not affect cell survival of neither P3 nor T98G cells. However, while BTZ treatment reduced P3 cell survival by 50% and could not be further augmented by addition of MRT68921 or VPS34-IN1, the cytotoxic efficacy of BTZ was annihilated by *ATG5* and *ATG7* depletion in P3 cells, which lacked LC3A/B-II and failed to accumulate autophagosomes compared BTZ treated controls (*P* < 0.0001 respectively, Fig. 5E). Neither *ATG5* nor ATG7 depletion could rescue T98G cells from BTZ induced cytotoxicity (Supplementary Fig. 2C). Addition of chloroquine enhanced BTZ cytotoxicity in P3 wild type cells. Clonogenic survival of *ATG5^-/-^* and ATG7 shRNA cells in response to TMZ treatment was not affected, nor when ULK1/2 or VPS34 was blocked with MRT68921 or VPS34-IN1 autophagy inhibitors in P3 cells (Fig. 5E). In contrast, TMZ effect was marginally improved by addition of VPS34-IN1 in T98G cells (*P* <0.05, Supplementary Fig. 2C). However, survival was augmented in both P3 *ATG5^-/-^* and ATG7 knockdown cells (*P* < 0.0001 and *P* < 0.01 respectively, Fig. 5E) and in only T98G *ATG5^-/-^* cells (*P* < 0.0001, Supplementary Fig. 2C) in response to BTZ+TMZ treatment relative to controls. The BTZ cytotoxicity is diminished in both *ATG5* and ATG7 depleted P3 cells while in T98G cells, the BTZ effect was only apparent in *ATG5^-/-^* ablated T98G cells treated with combination BTZ+TMZ. In contrast, attenuated LC3A/B-II formation through inhibiting ULK1/2 or VPS34 complexes was also reflected in differential killing efficacy of P3 and T98G GBM cells, as was treatment with BTZ alone.

### *Autophagy abrogation increased MAP1LC3B/p62 secretion in ATG5^-/-^ P3 cells*, as well as after BTZ and BTZ+TMZ treatment

Untreated *ATG5^-/-^* P3 cells accumulated more autophagosomes than autolysosomes (*P*<0.01) also compared to BTZ treated cells (*P*<0.05, Fig. 5D), yet lysates contained undetectable LC3A/B-II protein. To investigate whether these autophagosomes might have been expelled to the extracellular matrix, we investigated the cellular secretome during nutrient-deprivation by liquid chromatography mass spectrometry. In total 3720 proteins were identified, where 2735 proteins were used for quantification. Hierarchical clustering from 164 statistically significantly secreted proteins identified 7 distinct biological processes involved in proteasome function, lysosomal metabolic processes, protein degradation and autophagy (Supplementary Fig. 3). P3 *ATG5^-/-^* cells or P3 wild type cells treated with BTZ or BTZ+TMZ combination showed similar clustering in secretory autophagy and mitophagy related proteins compared to supernatants from control and TMZ treated cells that clustered more closely together (Supplementary Fig. 3A). More MAP1LC3B and p62(SQSTM1) were secreted into the supernatants in *ATG5^-/-^* P3 cells or when autophagy flux was abrogated with BTZ or BTZ+TMZ treatment compared to TMZ only or untreated controls (*P* <0.001 and *P*=0.007, Supplementary Fig. 3B). Taken together, our data indicate that BTZ kills GBM cells by inhibiting autophagic flux, prohibiting autophagosome-lysosome fusion and subsequent degradation of cargo.

### BTZ induces DNA damage signaling and cell cycle arrest in TMZ treated GBM cells

Since clonogenic survival after TMZ treatment appeared to be sustained by autophagic flux, and BTZ alone or in combination with TMZ promoted GBM cell death, we sought to investigate the crosstalk between abrogated autophagic flux and cell death, as well as characterize the mechanism of death. First, we examined effects of the treatments on DNA damage and perturbation of cell cycle kinetics. Thus, we investigated the phosphorylation status of the DNA damage sensors ATM, H2AX and Chk2 in chemoresistant GBM cells (P3 and T98G) in response to BTZ and TMZ mono- or combination therapy. At the IC_50_ dose of either TMZ or BTZ alone, the phosphorylation of serine and threonine residues of ATM (Ser1981), H2AX (Ser139) and Chk2 (Thr68) was consistent with initiation of DNA damage signaling cascade at 48h in P3 cells (Fig. 6A and 6B). These changes in protein phosphorylation and downstream mediators were delayed in response to TMZ monotherapy compared to BTZ monotherapy (Fig. 6A and 6B). BTZ monotherapy induced 3-fold p53 and p21 expression within 24h compared to TMZ monotherapy, where increases in phosphorylated ATM, H2AX, and p53 (Ser15), as well as the downstream genes Chk2 (5-fold) and p21 was 3-fold upregulated after 48h (Fig. 6A and 6B). Treatment of P3 cells with a combination of TMZ and BTZ led to 2-fold increased phosphorylation of ATM, H2AX and p53 after 48 h, but with half the TMZ IC_50_ dose (Fig. 6C), implying augmented DNA damage response. Similar responses were confirmed in T98G cells treated with TMZ, where ATM (Ser1981) and H2AX (Ser139) together with phospho-Chk2 (Thr68) were strongly expressed at 48h and were maintained up to 120h (Supplementary Fig. 4A–4C). These results suggest that BTZ potentiates the activation of cell cycle checkpoints and accumulation of proteins involved in DNA damage signaling.

**Figure 6.**
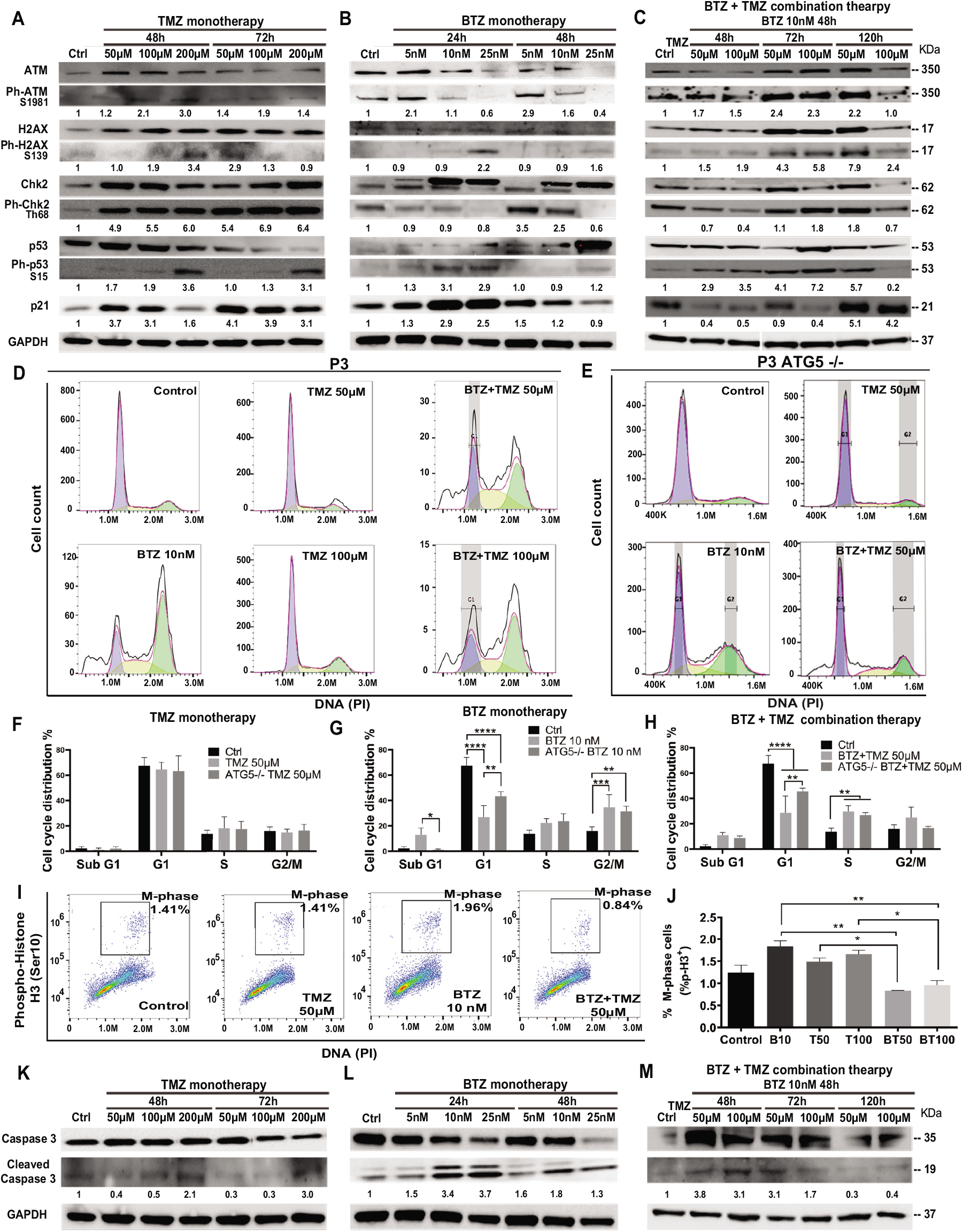
BTZ pretreatment induces DNA damage signaling and G2/M phase cell cycle arrest in TMZ treated patient derived GBM cells. Proteins in DNA damage signaling cascade including total and phospho-proteins from lysates (20 μg) from P3 GBM cells treated with (A) TMZ monotherapy, dose escalation at 48 and 72h, (B) BTZ monotherapy, dose escalation at 24 and 48h and (C) combination treatment with BTZ 10nM 48h and TMZ at indicated times. (D-E) DNA histograms showing cell cycle distribution of propidium iodide (PI) stained control and *ATG5^-/-^* P3 cells, before and after respective treatments. Quantification of cellular proportions (%) in respective cell cycle phases after (F) TMZ monotherapy, (G) BTZ monotherapy and (H) BTZ+TMZ therapy. (I) Dot plots of P3 cells in M-phase (phospho-histone H3 positive cells) after the respective treatments; (J) Quantification of experiments shown in panel I, showing % of P3 cells in M-phase of the cell cycle as indicated. Western blot analysis of total and cleaved caspase 3 in proteins lysates from P3 GBM cells treated with (K) TMZ monotherapy, (L) BTZ monotherapy and (M) combination treatment with BTZ+TMZ, at indicated times. GAPDH was used as a loading control and for normalizing densitometry measurements of the blots. *P < 0.05, ** P < 0.01, *** P < 0.001 and **** P < 0.0001.

Cell cycle kinetics were examined in P3 cells to determine whether cell cycle checkpoints were executed in response to the three treatment regimens. As anticipated TMZ alone did not substantially alter cell cycle checkpoints nor alter apoptosis threshold in TMZ resistant P3 cells (Fig. 6D and 6F). BTZ 10nM treatment however, led to a significantly reduced cellular portions entering G1 compared to untreated control P3 cells (*P* < 0.0001), with a concomitant increase in G2/M phase arrest (*P* < 0.001) as well as increased apoptotic proportions in hypodiploid sub-G1 phase (*P* < 0.05), Fig. 6D and 6G). *ATG5^-/-^* P3 cells’ cell cycle progression was not altered by TMZ treatment (Fig. 6E and 6F). However, *ATG5^-/-^* P3 cells increased cell proportions entering G1 (*P* < 0.01) and diminished apoptotic fraction in sub-G1 (*P* < 0.05) compared BTZ treated control cells, essentially reversing the BTZ cytotoxic effect (Fig. 6D-E and 6G). BTZ+TMZ combination treatment also significantly reduced cell fractions in G1 phase compared to untreated P3 control cells (*P* < 0.0001), and *ATG5^-/-^* ablation in P3 cells partially re-established the cell proportions entering G1-phase (*P* < 0.01, Fig. 6D-E; 6H). ATG7shRNA knockdown modestly prevented P3 cell cycle progression to G1 (*P* < 0.0001) and increased G2/M phase arrest after TMZ (*P* < 0.001, Supplementary Fig. 5A and 5B). Importantly, ATG7shRNA knockdown modestly reversed BTZ induced block in P3 cells’ G1 phase transition (*P* < 0.05, Supplementary Fig. 5A and 5C) while ATG7shRNA knockdown in P3 cells had less effect than BTZ+TMZ treatment (Supplementary Fig. 5A and 5D). Both *ATG5^-/-^* and ATG7shRNA knockdown reduced T98G cell fractions entering G1 cell cycle phase after BTZ treatment (*P* < 0.001 respectively, Supplementary Fig. 5F, 5J, 5H and 5L) and increased G2/M arrest compared to untreated cells (*P* < 0.0001, respectively). Neither *ATG5^-/-^* nor ATG7shRNA knockdown in T98G cell had profound effect on responses to BTZ+TMZ combination treatment (Supplementary Fig. 5F, 5J, 5I and 5M).

This prominent effect of the combination treatment on cell cycle progression was also apparent when examining cell cycle kinetics at the molecular level. Cell fractions expressing phosphorylated histone H3 (Ser10), which is unique to the mitosis-phase of cycling somatic cells were significantly reduced by the combination therapy in P3 cells, compared to TMZ (*P* < 0.05) or BTZ (*P* < 0.01) monotherapy (P3 cells, Fig. 6I–6J; and in T98G, Supplementary Fig. 4E). Finally, BTZ at lower doses alone or in combination with TMZ induced 4-fold increased caspase 3 cleavage in P3 cells (Fig. 6L–6M), while caspase 3 was not cleaved following TMZ monotherapy (Fig. 6K). Only high dose BTZ 25nM monotherapy or combination BTZ+TMZ induced cleavage of caspase 3/8 in T98G cells (Supplementary Fig. 4G-H). Taken together, these results indicated that BTZ pretreatment sensitized GBM cells to TMZ-induced DNA damage and programmed cell death. Perturbing the autophagy machinery with CRISPR-Cas9 *ATG5^-/-^* and ATG7 shRNA knockdown eliminated the cytotoxic efficacy of BTZ containing treatments in autophagy dependent P3 cells, but not T98G cells.

Abrogated autophagy flux is postulated to decrease the apoptosis threshold through mechanisms involving the autophagy-regulating transcription factor FOXO3a and p53-upregulated modulator of apoptosis (PUMA). Chemoresistent tumor cells with high autophagy turnover inhibit transcriptional activation of FOXO3a and PUMA^21,45,46^. Thus, we investigated the implication of this mechanisms in BTZ mediated blockade of autophagic flux in sensitizing GBM to TMZ. Although no significant differences in PUMA mRNA levels were induced by treatment (Supplementary Fig. 5E), FOXO3a mRNA levels were upregulated by approximately 2-fold in BTZ mono- and BTZ+TMZ combination treated P3 GBM cells respectively (*P* < 0.05) compared to TMZ treatment alone.

## Discussion

In this work we report that TMZ resistant tumors upregulate autophagic flux, a metabolic process which cancer cells usurp as an adaptive survival mechanism in order to sustain themselves through recycled amino acids and other metabolic precursors from the degraded cytoplasmic components^17–19^, especially during conditions of stress, such as chemotherapy. P3 cells had high basal autophagy flux under steady state and during TMZ treatment, as denoted by low levels LC3A/B-II and p62(SQSTM1), equal measures of autophagosome and autolysosomes and corroborated by increased degradation of long-lived proteins. TMZ treatment increased the fraction of phospho-histone-H3 positive cells in M-phase cell cycle, as well as limited apoptotic fractions compared to control cells. These findings are consistent with increased autophagic flux as a pro-survival mechanism. Although both T98G and P3 cells had mutations in *PTEN*, P3 cells harbored additional mutations in *PIK3R2, MAP3K1* and *STAT3* genes that potentially increase downstream PI3K activity and influence autophagy dependence^47^. This is important because it has previously been shown that autophagy dependent tumors respond well to proteasome inhibition^48^. Further supporting the findings that drug synergy between inhibitors of autophagic flux and anti-neoplastic drugs occur more frequently in autophagy-dependent tumors we demonstrated that BTZ at 15nM was synergistic with all doses of TMZ, augmenting both P3 and T98G GBM cell killing. T98G cells were however, less prone to LC3A/B lipidation and cell death by BTZ alone or CQ compared to P3 cells. Pertubation of core autophagy machinery proteins had less robust effects on T98G cells potentially indicating a less profound dependence on this metabolic process. Specific mutations in critical pathways influence whether a particular cancer type becomes autophagy addicted. Genomic heterogeneity might explain conflicting data regarding efficacy of autophagy inhibitors on different cell types where the same drug combination might conceivably be anti-agonistic in autophagy-independent tumor types^48,49^.

Blocking the 26S proteasome with BTZ alone or in combination treatment with TMZ abrogated autophagic flux as evidenced by accumulation of LC3A/B-II that was not further increased in the presence of lysosomal inhibitor BafA1, which prevents autophagosome-lysosome fusion^39^. Concomitantly increased p62(SQSTM1) protein in lysates and intracellular compartments from P3 cells, as well as relative numbers of autophagosomes and autolysosomes on electron micrographs confirmed accumulation of autophagosomes. The accumulation of autophagosomes was due to lack of fusion with lysosomes with subsequent attenuated degradation of cargo. In line with this, STX17 did not colocalize with LC3A/B and the turnover of long-lived proteins attenuated after BTZ+TMZ combination treatment. Through multiple approaches with both static and dynamic consensus methods^14,50^, including LLPD assays, electron and confocal microscopy to assess the intracellular dynamics of autophagic vesicles, and quantification of the autophagic compartments, we were able to demonstrate consistently that the ratio of autophagosomes to degradative autolysosomes was significantly increased under BTZ+TMZ combination treatment.

BTZ has been contradictorily proposed to induce autophagy^40,41^. Since induction of LC3A/B-II on immunoblots indicates increased autophagosomes, most studies unilaterally concluded this indicated autophagy induction. However, without investigating whether cargo degradation takes place, one cannot conclude solely based on LC3A/B lipidation since accumulation of autophagosomes can indeed be due to increased flux or decreased cargo degradation. Therefore, it is imperative to quantify levels of LC3A/B-II induced by the given treatment and in the presence and absence of the lysosomal protease inhibitor BafA1^44^ and corroborate findings with gold standard dynamic flux assays of long-lived protein degradation^19^. Our results are in line with other studies demonstrating that BTZ blocks the catabolic process of autophagy^42,43^. Treatment of ovarian cancer cells with BTZ, as well as another proteasome inhibitor MG-132, was shown to block autophagic flux by increasing GFP-LC3 puncta, LC3A/B-II and p62(SQSTM1) protein levels and the number of autophagosomes^42^. In another study, BTZ treated cells showed higher LC3A/B-II and p62(SQSTM1) levels and inhibition of the turnover of long-lived proteins in serum starved and rapamycin treated MCF-7 cells, conclusive with blockade of the catabolic process of autophagy^43^. In agreement with these studies, we determined that BTZ sensitized GBM cells to TMZ by abrogating the autophagic flux at a late stage of the multi-stage process. BTZ treatment strongly induced lipidation of LC3A/B and autophagosome accumulation, similar to treatment with the lysosomal inhibitors BafA1 and CQ. In contrast, depletion of ATG5 or ATG7, proteins that are critical for cargo recruitment and elongation of the phagophore^51^, both abolished lipidated LC3A/B and ATG5^-/-^ rescinded the autophagosome formation in response to BTZ treatment and rescued the cells from BTZ induced death, confirming that BTZ effect was *ATG5* dependent. However, untreated *ATG5^-/-^* cells did contain more autophagosomes compared to BTZ treated *ATG5^-/-^* cells, supported by a recent study that demonstrated that cells lacking ATG7 still form autophagosomes through an alternative pathway^13^. LC-MS proteomics of the cellular secretome revealed greater levels of MAP1LC3B and p62(SQSTM1) in supernatants from the untreated *ATG5* cells. Instead of these autophagosomes fusing with lysosomes to degrade cargo, they might have fused directly with the plasma membrane to expel autophagic content to the extracellular space^52^. Hence MAP1LC3B and p62(SQSTM1) were detected in the secretome but not cellular lysate in the ATG5^-/-^ cells. Most importantly, this failure to increase autophagosome formation under BTZ treatment in *ATG5* and ATG7 knockout P3 and T98G cells rescued them from death by treatment with BTZ alone or in combination with BTZ+TMZ, indicating that cell death induced by BTZ is robustly dependent on an ATG5 functional autophagy machinery. In concurrence, *ATG5^-/-^* and ATG7 shRNA knockdown reversed the G1 checkpoint arrest, and sub-G1 apoptotic fractions that were induced by BTZ treatment, particularly in the autophagy dependent P3 cells. Notwithstanding the short-term analysis of the cell cycle kinetics was robustly consistent with the findings from longterm colony formation analyses.

In line with this, pharmacological perturbance of the early autophagy components ULK1 and VPS34 with inhibitors MRT68921 and VPS34-IN1, respectively, also abolished lipidated LC3A/B protein, although this had less impact on BTZ induced cell death. Several explanations can account for the differences in autophagic flux and abolished cell death observed in ATG5 and ATG7 depleted cells versus cell treated with inhibitors of the upstream autophagy kinases ULK1/2 and VPS34. The observed block of autophagic flux can be caused by effects of VPS34 on endocytosis. It is interesting to note that ATG5 and ATG7 depletion also reduced cell survival in control cells, although the effects were different from cells treated with ULK1 and VPS34 inhibitors upon co-treatment with BTZ. This might be explained by our finding that ULK1 total and phosphorylated protein levels are diminished in BTZ treated cells, which may cause GBM cell death. A variable ability of some proteasome inhibitors to block macrophagy, but not chaperon-induced autophagy, may also play a critical role in clearance of aggregated proteins, or damaged organelles in some cell types^53–55^. BTZ is a reversible inhibitor and we previously showed that the activity of the 20S proteasome returns to baseline within 72h^56,57^. Thus, the timespan of 120 hours encompassing the BTZ+TMZ combination treatment might have allowed partial recovery of autophagic flux.

Another mechanism of TMZ resistance in GBM patients is represented by unmethylated *MGMT* promoter tumors^58^, where the enzyme and proficiently repairs the chemotherapy-induced DNA damage but the cells maintain a high apoptosis threshold. We recently showed that BTZ treatment depletes MGMT protein and mRNA and sensitizes GBM cells to TMZ chemotherapy^59^. Abrogated autophagic flux would thus represent an additional mechanism by which BTZ kills GBM permitting more rapid DNA damage response by phosphorylation dependent activation of ATM, H2AX, and p53 as well as downstream Chk2 and p21. BTZ+TMZ combination therapy retained elevated phosphorylation of ATM and H2AX induced by BTZ pretreatment, reduced portion of cells in G1 phase with a concomitant increase in G2/M phase and increased the apoptotic hypodiploid sub G1-fraction. Cell fractions expressing phosphorylated histone H3 (Ser10) were reduced by the combination BTZ+TMZ indicating sensitization to TMZ. BTZ abrogated autophagic flux lowered the apoptosis threshold, indicated by increased FOXO3a compared to TMZ treatment, potentially inducing PUMA and apoptosis. These findings are corroborated by a body of work elucidating the molecular crosstalk between autophagy inhibition and apoptosis^46,60–62^.

In conclusion, BTZ augmented cell death of TMZ resistant tumor cells by accumulating autophagosomes as a result of abrogated autophagic flux (Fig. 7), promoting ATM/Chk2 mediated DNA damage signaling and activating p53-mediated G2/M cell cycle arrest^63^. As we recently demonstrated^59^, BTZ also depleted MGMT protein and mRNA, an additional mechanism that synergized with TMZ to effectively kill GBM cells by enhanced programmed cell death. We anticipate to uncover similar mechanisms of efficacy in the patients undergoing treatment in our phase II clinical trial, NCT03643549 (www.clinicaltrials.gov).

## Material and methods

### Isolation of primary glioma cells

P3 patient-derived GBM cells were expanded from biopsies obtained from patients undergoing tumor resection at the neurosurgery department (Haukeland University Hospital; Bergen, Norway), following biobank (REK Vest 013.09/20879) and project (REK2018/71) approval from the regional ethical committee, the Norwegian Data Protection Agency and patient informed consent. These cells, as well as long-term established U87, A172, and T98G cell lines (ATCC, Manassas, VA, USA) and HF66 (Henry Ford Institute, Detroit, MI, USA) were propagated in Dulbecco’s modified eagle medium (DMEM, Sigma-Aldrich; St. Louis, MO, USA) supplemented with 10% fetal bovine serum, non-essential amino acids, 100 U/mL penicillin/streptomycin and 400 μM L-glutamine (complete medium; all Cambrex; East Rutherford, NJ, USA) at 37°C in a humidified atmosphere of 5% CO_2_.

### Drugs

Temozolomide (2706/50, Tocris Bioscience, Bristol, UK) was dissolved in dimethyl sulfoxide (DMSO and stored at −20°C. Bortezomib (Velcade^®^, 576415, Janssen, Norway) (Haukeland University Hospital pharmacy, Bergen, Norway) was dissolved in 0.9% v/v sodium chloride and stored at −80°C. ULK1 inhibitor (MRT68921, Cat. No. S7949, Selleckchem, USA) and VPS34 inhibitor (VPS34-IN1, Cat. No. S7980, Selleckchem, USA) were dissolved in DMSO and stored at −20°C. Bafilomycin A1 (Cat. No. SML1661) and Chloroquine diphosphate salt (Cat. No.C6628) were obtained from Sigma-Aldrich. Chloroquine was dissolved in water and stored at −20°C.

### Clonogenic survival assay

Cells were seeded at plating efficiency density^64^ (1000 cells/well) in 6-well plates and exposed to temozolomide (72 h: 6–500 μM) or BTZ (48 h: 1-25nM) or 1h incubation with MRT68921 (1μM) or VPS34-IN1 (1 μM), monotherapy or in combination, followed by further observation for 14 days. Colonies were stained with crystal violet and counted as previously described ^65,64^. IC_50_ were calculated using Prism software version 6.07 (GraphPad; La Jolla, CA, USA). Compusyn software was used for drug combination synergy calculation following the Chou-Talalay method^66^. All experiments were performed in triplicate and repeated at least 3 independent times.

### Western blot analysis

Cells were lysed in Kinexus lysis buffer: 20 mM MOPS, 5 mM EDTA, 2 mM EGTA, 30 mM NaF, 0.5% Triton X, 1 mM PMSF, pH 7.2, protease inhibitor (cocktail tablet, Roche; Basel, Switzerland), and phosphatase inhibitor (cocktail tablet, Roche). Samples (10-20 μg) were run on SDS/PAGE with NuPAGE precast 4-12% gradient gels (Invitrogen; Carlsbad, CA, USA) and blots were incubated overnight at 4°C with primary antibodies (Supplementary Table 1) followed by incubation for 1.5 h at room temperature (RT) with a species specific secondary HRP-conjugated antibody diluted 1:10000. Chemiluminescence detection was performed with Super Signal West Femto Maximum Sensitivity Substrate (Thermo Fisher Scientific, Bremen, Germany) on the LAS-3000 (Fujifilm Medical Systems Inc.; Stamford, Connecticut, USA). Relative protein expression levels were normalized to GAPDH and quantified using Image J software (NIH; Bethesda, MD, USA).

### Cell cycle distribution and kinetics

P3 and T98G cells with and without CRISPR-Cas9 *ATG5^-/-^* and ATG7shRNA knockdown were treated with TMZ (72 h: 50 μM and 100 μM for P3; 125 μM and 250 μM for T98G) or BTZ (48 h: 10nM) monotherapy or in combination. Briefly, for cell cycle distribution analysis, cells were harvested post-treatment, fixed in ice-cold ethanol and stained with 50 μg/mL propidium iodide (PI) or phospho-histone H3 (Ser10) (Supplementary Table 1), as previously described^65^. Data were acquired on a FACS Accuri™ flow cytometer (BDbiosciences; Franklin Lakes, NJ, USA), and analyzed with FlowJo^®^ software (Treestar Inc; Ashland, OR, USA). The Dean-Jett-Fox algorithm with synchronized peaks was used to analyse the DNA histograms of single cells from three independent experiments.

### Autophagic flux assay

The autophagic flux assay was carried out using the Premo™ Autophagy Tandem Sensor RFP-GFP-LC3 Kit (P36239, Molecular Probes, Thermo Fisher Scientific, Bremen, Germany), following the manufacturer’s instructions. Briefly, 2.5 × 10^4^ P3 cells were cultured on coverslips in 24-well plates overnight, incubated with 10 μL of BacMam reagent for 24h, and then treated according to schedule with BTZ (10nM), TMZ (50 and 100μM) or in combination. 50μM Chloroquine (CQ) was added 24h before end of the BTZ, TMZ or combination treatments. Cells were washed, slides mounted with Prolong Gold DAPI (P36931, Molecular Probes) and confocal images acquired on a Leica T SC SP5 3X (Leica, Wetzlar, Germany). The number of LC3B-positive autophagosomes and autolysosomes in merged images was quantified in at least 10 cells from each treatment group.

### Long-lived protein degradation

P3 and T98G cells were seeded at density of 4 × 10^4^ cells/well in 24-well plates, and treated with BTZ, TMZ or in combination. To measure the degradation of long-lived proteins by autophagy, proteins were first labeled for 48 h with 0.25 μM Ci ml^-1^ L-^[U-14C]^ VALINE (Perkin Elmer, Waltham, MA, USA) in GIBCO-RPMI 1640 medium (Invitrogen, Carlsbad, CA, USA) containing 10% FBS, washed in warm PBS and then chased for 18 h without radioactivity in DMEM containing 10% FBS and 10 mM valine (Sigma-Aldrich), to allow degradation of short-lived proteins as previously described^67^. 50 μM CQ was added 24h before cell harvest. After 24h, the supernatant was collected, 50% TCA was added and proteins precipitated over-night at 4 °C. The cells were lysed with 0.2M KOH over-night at 4°C. The supernatant was centrifuged and transferred to a new tube, the precipitate dissolved and moved to the same sample as the cell lysate. Both supernatant and lysate were moved to separate counting vials and mixed with 3 mL scintillation fluid (Ultima Gold #6013321, Perkin Elmer). ^14^C levels were measured in each sample using a Packard Liquid Scintillation Analyzer. The percent degradation was calculated by comparing the amount of ^14^C in the supernatant to the total ^14^C levels (supernatant and lysate). The autophagic flux was calculated by subtracting the percent degradation of the CQ treated sample from the untreated sample, for each culture medium. All samples were related to the control.

### Generation of ATG5 knockout cells using CRISPR-Cas9 genome editing

Human *ATG5* knockout P3 and T98G GBM cell lines were generated by using LentiCRISPRv2^68^ (purchased from Addgene, plasmid# 99573). Plasmids were transfected into 293T cells using BBS/CaCl_2_ to produce lentivirus. The viral supernatant was harvested at 48h post-transfection, filtered through 0.45-μm filters (Millipore). Infection of P3 and T98G cells was performed by centrifugation of cells with virus at 2225 rpm for 90 minutes at room temperature in the presence of 10 μg/mL polybrene (Sigma-Aldrich). Enrichment selection using puromycin (1 μg/mL) (Sigma-Aldrich) was done and collected for gene knockout assessment by western blotting.

### Atg7 RNA interference

Human ATG7 shRNA-GFP (5’-CAGTGGATCTAAATCTCAAACTGAT-3’) and scrambled control shRNA-GFP (5’-GGGTGAACTCACGTCAGAA −3’) lentiviral plasmids were purchased from Applied Biological Materials Inc. (Richmond, BC, Canada). Plasmids were transfected into 293T cells using BBS/CaCl_2_ to produce lentivirus. Infection of P3 and T98G cells was performed by centrifugation of cells with virus at 2225 rpm for 90 minutes at room temperature in the presence of 10 μg/mL polybrene (Sigma-Aldrich). Enrichment selection using puromycin (1 μg/mL) (Sigma-Aldrich), followed by FAC-Sorting on Sony SH800 (Sony Biotechnology, San Jose, CA, USA) yielded stably expressing shRNA cells.

### Quantification of autophagic flux by CYTO-ID Flow Cytometry Assay

Autophagic flux was induced by serum starvation through 3h culture in Earl’s balanced salt solution (EBSS) supplemented with 0.1% BSA). Inhibition of autophagic flux by blocking fusion between autophagosomes and lysosomes was induced by treatment with CQ (50μM) for 24h, final concentration 50μM for 3h). Cells were treated with BTZ 10nM for 48h, with and without CQ 50μM for 24h. CYTO-ID Autophagy detection kit 2.0 (ENZ-KIT175-0050, Enzo Life Sciences, Farmingdale, NY, USA) was used and to monitor the autophagic flux at the single-cell level as previously described^61^ and in accordance with manufacturer’s instructions. The dynamic autophagosome generation and clearance was assessed by live cell flow cytometry using BD Accuri C6 flow cytometer (BD Biosciences). Median fluorescence intensities (FL2-A channel) of 5000 cells are given. The dye of the Cyto-ID assay incorporates into autophagic vacuoles (pre-autophagosomes, autophagosomes, and autolysosomes (autophagolysosomes), and thus allows single cell measurement of autophagic flux in lysosome inhibited live cells.

### Immunocytochemistry and confocal microscopy

Cells (4×10^4^ cells/well) were plated on coverslips in 24-well plates and subsequently treated with the various drug combinations as indicated above for 48 or 72h. For LC3A/B and p62(SQSTM1) and STX17 (Supplementary Table 1) staining, live cells were fixed with 3.7% formalin for 15 min at RT, stained with primary antibodies, followed by incubation with goat anti-rabbit-Alexa Fluor^®^555 and goat anti-mouse-Alexa Fluor^®^488 (Supplementary Table 1). Slides were mounted with Prolong Gold DAPI and images acquired on Leica TSC SP8 STED 3X (Leica).

### Transmission electron microscopy

Treated or control cells were fixed overnight in 2% glutaraldehyde in cold medium and thereafter rinsed in 0.1 M Na-cacodylate buffer prior to post fixation in 1% osmium (OsO_4_) in Na-cacodylate buffer for 1 h. Cells were washed again in 0.1 M Na-cacodylate buffer dehydrated in 30% ethanol for 20 min on ice, 50% ethanol for 20 min at 4°C and finally, 70% ethanol overnight at RT. The second dehydration cycle was the following: 96% ethanol for 30 min at RT, 100% ethanol for 30 min (×2), and propyleneoxide for 20 min. The sample was mixed in propylene oxide/agar 100 resin in a 1:1 ratio overnight at RT, further hardened overnight at 60°C prior to sectioning to 45 - 50 nm. Imaging was performed on a Jeol JEM-1230 (Tokyo, Japan,) at 80 kV using a gatan camera. The number of autophagosomes and autolysosomes were counted from at least 10 cells from 2-3 independent experiments by two independent investigators.

### RT-qPCR

RNA from P3 cells treated with BTZ alone, TMZ alone or combination with BTZ and TMZ were isolated using Qiagen RNeasy kit following the manufacturer’s instructions. cDNA was synthesized using the iScript cDNA Synthesis kit (BIO-RAD; Hercules, California, USA) according to the manufacturer’s instructions. iQ SYBR Green from the Supermix kit (BIO-RAD) was used to detect amplified produce in the PCR reaction mixture. The reaction was run on a Roche light cycler (LC480, Roche; Indianapolis, IN, USA) for 40 cycles. The primer sequences were as follows: ULK1 forward 5’-GGACACCATCAGGCTCTTCC-3’ and reverse 5’-GAAGCCGAAGTCAGCGATCT-3’, FOXO3a forward 5’-GCGACAGCAACAGCTCTGCC-3’ and reverse 5’-GGGCTTTTCCGCTCTTCCCCC-3’, PUMA forward 5’-TCAACGCACAGTACGAGCG-3’ and reverse 5’-AAGGGCAGGAGTCCCATGAT-3’ and internal control 18S forward 5’-CGGCTACCACATCCAAGGAA-3’ and reverse 5’-GCTGGAATTACCGCGGCT-3’. Target transcripts were normalized to 18S and analyzed using the comparative CT (ΔΔCT) method.

### Liquid chromatography mass spectrometry

The protein secretome in conditioned media of treated cells was analysed by Electrospray ionization LC-MS. Samples from two independent experiments were run on an Ultimate 3000 RSLC system (Thermo Scientific, Sunnyvale, California, USA) connected online to a Q-Excative HF mass spectrometer (Thermo Scientific, Bremen, Germany) equipped with EASY-spray nano-electrospray ion source source (Thermo Scientific). MS spectra (from m/z 375-1500) were acquired in the Orbitrap with resolution R = 60000 at m/z 200, automatic gain control (AGC) target of 3e6 and a maximum injection time (IT) of 110ms. Complete methods are described in detail in supplementary methods.

### Mutational analysis of P3 and T98G GBM cells

Targeted sequencing of 360 cancer genes, was performed as previously described^70^. Briefly, native, genomic DNA isolated from cell lines was fragmented and Illumina DNA sequencing libraries were prepared. Libraries were then hybridized to custom RNA baits according to the Agilent SureSelect protocol. Paired-end, 75bp sequence reads were generated on an Illumina MiSeq instrument. Sequencing coverage for the targeted regions (average per bp) for the sample were 256x (P3) and 253x (T98G), supplementary Table 2.

Raw mutations were called by the MiSeq Local Run Manager software and annotation was performed applying Annovar. Called variants were restricted to those passing all default filters and further filters were: including only coding variants, excluding synonymous variants, variant allele frequency >0.1, population frequency <0.005 in 1K genomes project “all” and “eur”, population frequency <0.005 in esp6500.

### Statistical Analysis

When comparing more than two groups with one dependent variable, one-way ANOVA was used, while two-way ANOVA was used to analyze data with two or more dependent variables compared in two or more groups. Bonferroni or Tukey’s posthoc correction for multiple testing was used. Descriptive statistics were reported as mean ± standard error of the mean (SEM) unless otherwise stated. Two-sided *P*-values less than 0.05 were considered significant, (shown as * *P* < 0.05, ** *P* < 0.01, *** *P* < 0.001, and **** *P* < 0.0001). All graphs represent the mean ± standard error of the mean (S.E.M.) of at least 3 independent experiments. All statistical analyses were performed in Stata version 13.1 (Texas, USA) or GraphPad Prism v6.07 (La Jolla, CA, USA)

## Declarations

### Ethics approval and consent to participate

The regional ethical committee (REKVest) and The Norwegian Data Protection Agency approved the establishment of a brain tumor and blood biobank (REK Vest 013.09/20879) and issued project approval. Patients provided informed consent for use of their tissue samples for research purposes (REK2018/71).

## Author contribution

Conducted experiments and analyzed data: MAR, SS, AE, ML, EB, CB, SK

Provided reagents: AS

Writing the manuscript: MAR, MC

Revising the manuscript: all authors revised the manuscript

Designing research and acquired funding: MC

## Consent for publication

All authors contributed, read and consented to publication of the final version of the manuscript.

## Data availability

The mass spectrometry proteomics data have been deposited to the ProteomeXchange Consortium via the PRIDE^71^ partner repository with the dataset identifier PXD021828.

## Acknowledgements

We thank the staff at the neurosurgical department and surgical theaters at Haukeland University Hospital for providing tumor samples for the Brain Tumor Biobank. We are grateful to Endy Spriet, Hege Avsnes Dale at the Molecular Imaging center (MIC) UiB, for electron and confocal microscopy assistance. Flow cytometry was performed at the flow cytometry center at the University of Bergen.

## Abbreviations

GBM: Glioblastoma
TMZ: Temozolomide
BTZ: Bortezomib
DNA: Deoxyribonucleic acid
ATG5: Autophagy related 5
ATG7: Autophagy related 7
MAP1LC3A/B: Microtubule-associated proteins 1A/1B light chain 3B
p62/SQSTM1: Ubiquitin-binding protein p62/ Sequestosome-1
STX17: Syntaxin 17
ULK: Unc-51 like autophagy activating kinase
ATM: ATM serine/threonine kinase
Chk2: Checkpoint kinase 2
Gy: Gray
MGMT: O6-methylguanine DNA methyltransferase
PIK3C3: Phosphatidylinositol 3-kinase catalytic subunit type 3
VPS34: Vacuolar Protein Sorting-associated protein 34
SNARE: Soluble NSF attachment receptor
FOXO: Forkhead homeobox-type O
MEK: Mitogen-activated protein kinase kinase
ERK: Extracellular signal-regulated kinases
PI3K: Phosphoinositide 3-kinases
FDA: Food and Drug Administration
μM: Micromolar
nM: Nanomolar
IC50: Half maximal inhibitory concentration
Wt: Wild type
IDH1: Isocitrate dehydrogenase 1
SNV: Single nucleotide variant
TP53: Tumor protein 53
MAP3K1: Mitogen-activated protein kinase kinase kinase 1
PTEN: Phosphatase tensin homolog
PIK3R2: Phosphatidylinositol 3-kinase regulatory subunit beta
CQ: Chloroquine
BafA1: Bafilomycin A1
PE: Phosphatidylethanolamine
ATG12: Autophagy related 5
ATG16L1: Autophagy related 16 like 1
LLPD: Long-lived protein degradation
RFP: Red fluorescent protein
GFP: Green fluorescent protein
CRISPR: Clustered regularly interspaced short palindromic repeats
Cas9: CRISPR associated protein 9
shRNA: Small hairpin ribonucleic acid
H2AX: H2A histone family member X
Ser: Serine
Thr: Threonine
PUMA: p53-upregulated modulator of apoptosis
STAT3: Signal transducer and activator of transcription 3
LC-MS: Liquid chromatography-mass spectrometry
REK: Regional Committees for Medical and Health Research Ethics
ATCC: American Type Culture Collection
DMEM: Dulbecco’s modified eagle medium
DMSO: Dimethyl sulfoxide
PI: Propidium iodide

---

**Figure 7. Schematic summary of BTZ mediated inhibition of autophagic flux and its role in abrogating TMZ resistance in GBM cells.** A) Autophagosome formation requires the conjugation of cytosolic LC3-I to membrane bound LC3A/B-II and the reaction is catalyzed by autophagy related proteins ATG7 and ATG5. p62(SQSTM1) is bound to the cargo inside the autophagosome. Subsequent fusion of the autophagosomes (AP) to lysosomes results in autolysosome (AL) formation and degradation of the cargo (autophagic flux). Macromolecules generated from degraded contents of the autophagosome are recycled to the nutrient and energy cellular reservoirs, facilitating cancer cell survival during conditions of stress. To overcome the autophagy mediated cancer cell survival mechanisms, a number of autophagy inhibitors were tested, including upstream inhibition by knock down of ATG genes, and downstream inhibition by pharmacological agents like chloroquine, Bafilomycin A1 or proteasome inhibitors. B) TMZ chemotherapy sustains autophagic flux and underlies chemoresistance indicated as high clonogenic survival of GBM cells treated with TMZ. BTZ inhibits autophagic flux by (i) increasing autophagosomes seen on TEM, blocking lysosome-autophagosome fusion (ii) attenuated degradation of long-lived proteins(iii) accumulation of LC3B-II and p62 and reduced STX17 protein lysates, (iv) increased yellow puncta in Autophagy Tandem Sensor RFP-GFP-LC3B assay, and, all culminating in reduced clonogenic survival of GBM cells. Pretreatment of GBM cells with BTZ prior to TMZ treatment effectively inhibits autophagic flux and significantly reduced the clonogenic survival of GBM cells. Knock-out of *ATG5* using CRISPR-Cas9 and Knockdown of ATG7 using shRNA abrogates LC3B-II accumulation and upregulated p62 levels and decreased the number of autophagosomes after BTZ or BTZ+TMZ combination treatment, as more autolysosomes were also observed. This might indicate conventional inhibition of autophagic flux as well as sustained flux by alternative autophagy pathway in *ATG5^-/-^* and shATG7 GBM cells. While BTZ and BTZ+TMZ combination treatment reduced the clonal survival of TMZ resistant GBM cells, knockdown of *ATG5* or ATG7 rescued GBM cell survival under BTZ or BTZ+TMZ combination treatment indicating that blockade in autophagic flux overcame TMZ resistance and contributed to GBM cell death.

AP= Autophagy Flux, P=Phagophore, AP= Autophagosome, AL= Autolysosome, TEM= Transmission electron microscopy, WB= Western Blot, FM= Fluorescence Microscopy, LLPD= Long lived protein degradation assay, R= Red puncta and Y= Yellow Puncta

**Supplementary Figure 1.**
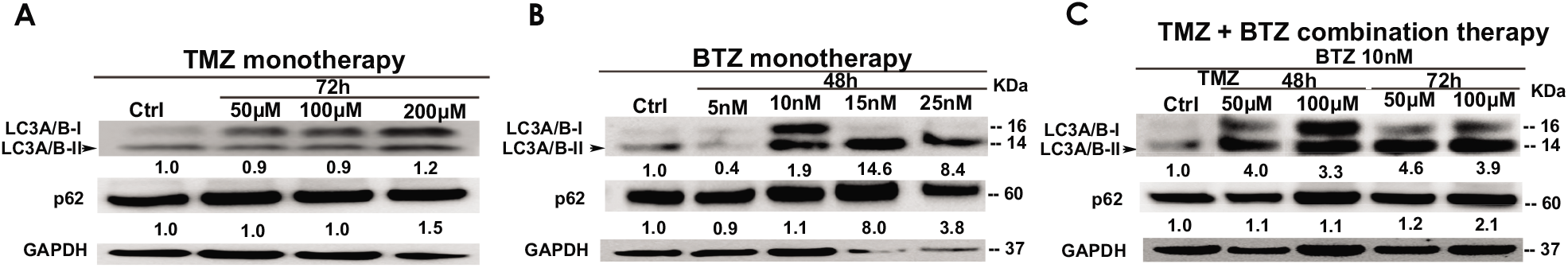

**Supplementary Figure 2.**
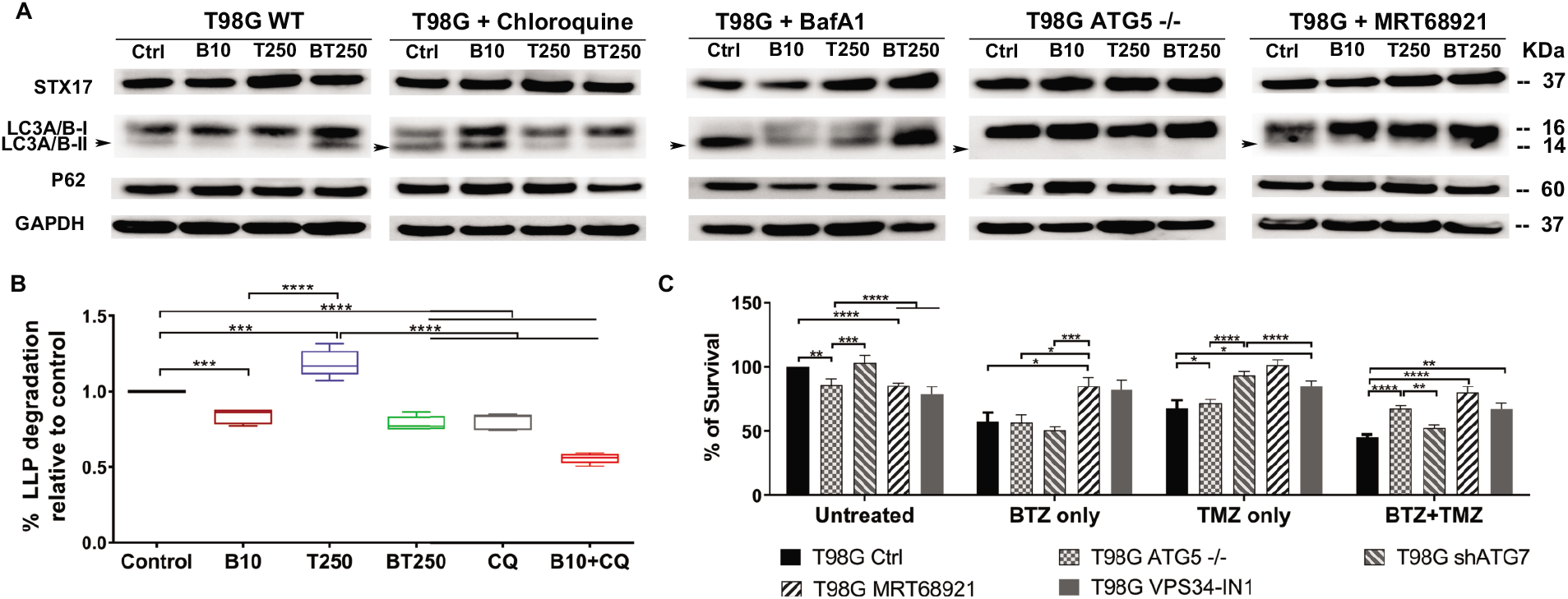

**Supplementary Figure 3.**
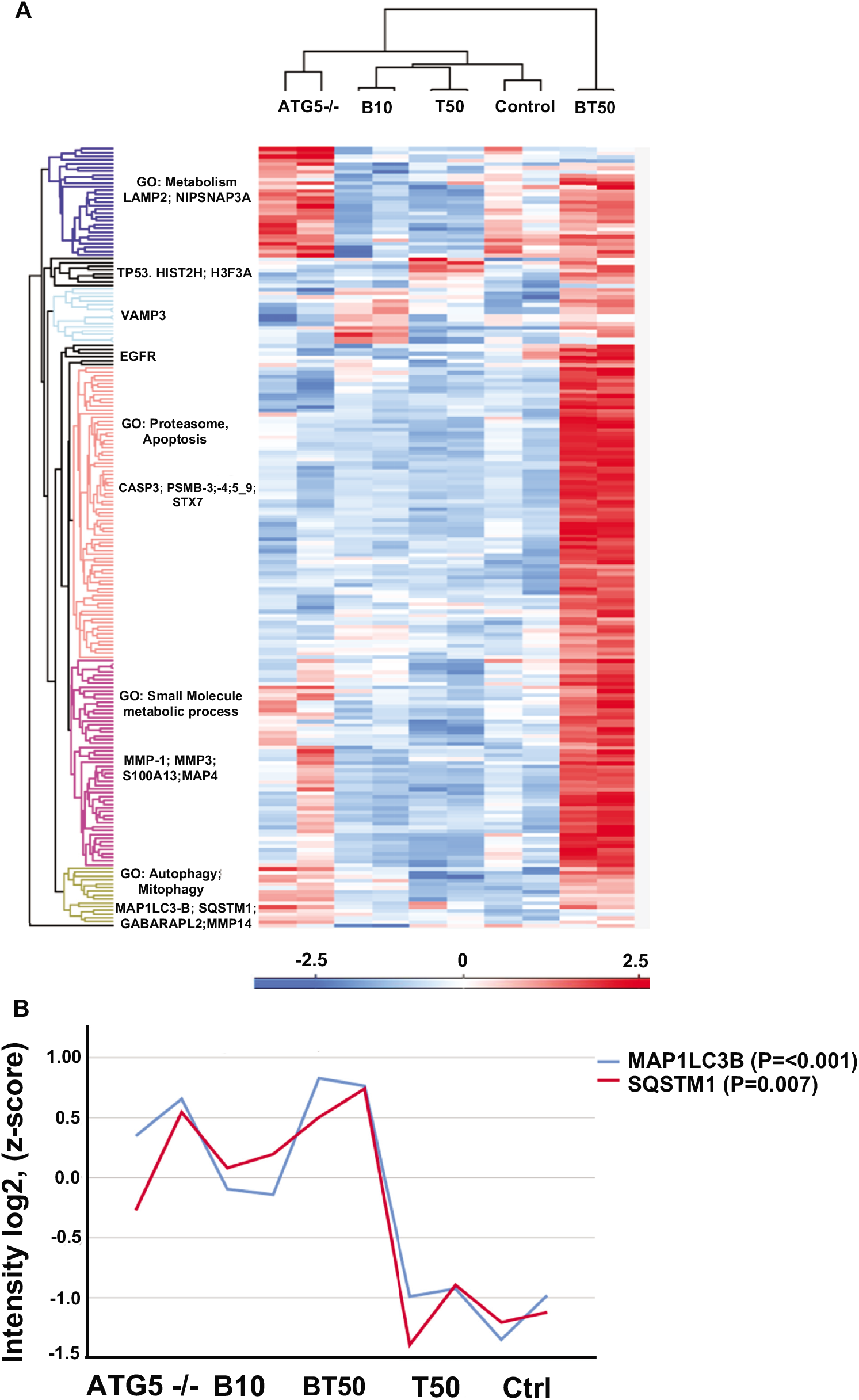

**Supplementary Figure 4.**
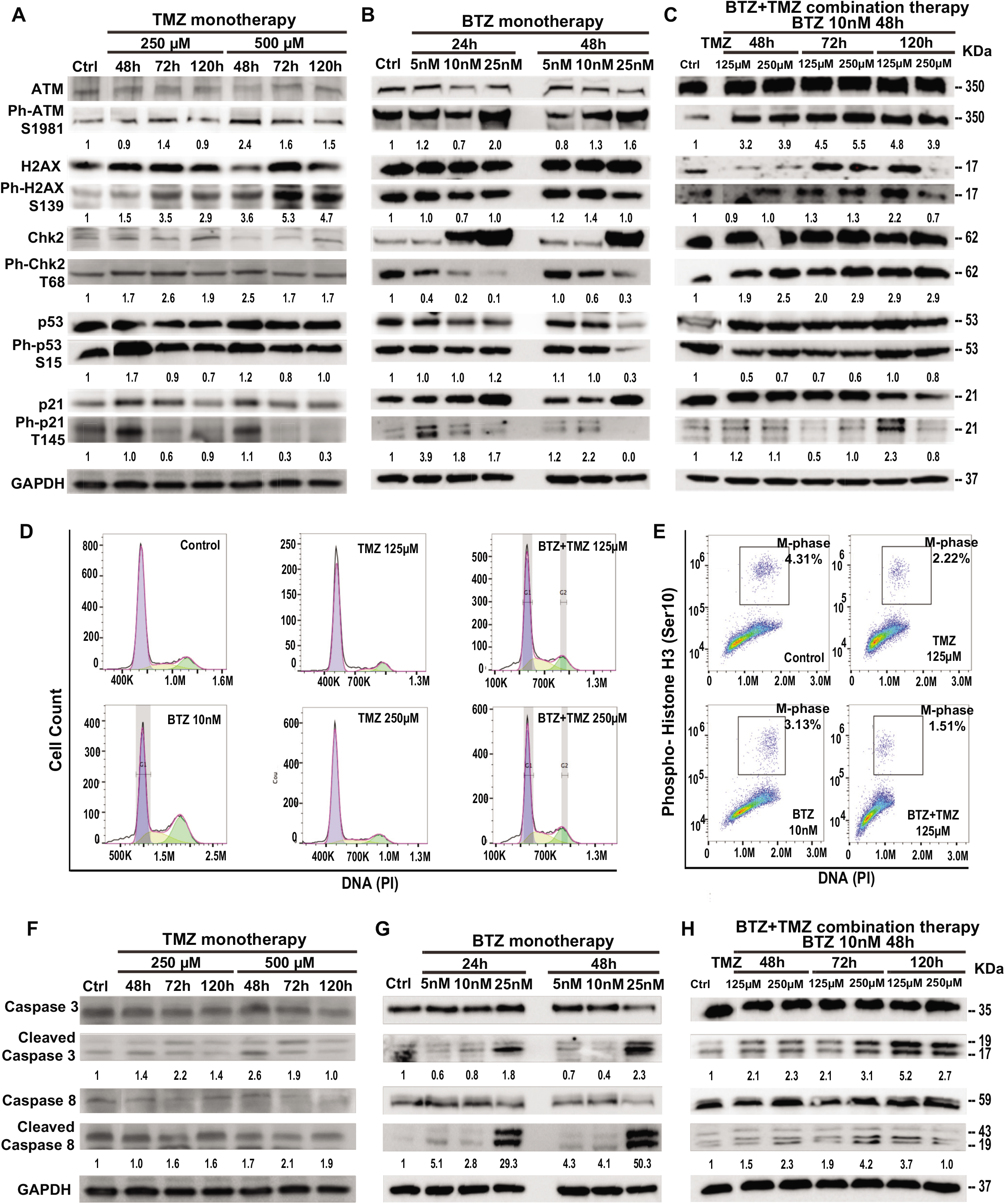

